# Coarse-Grained Model of Serial Dilution Dynamics in Synthetic Human Gut Microbiome

**DOI:** 10.1101/2024.01.23.576928

**Authors:** Tarun Mahajan, Sergei Maslov

**Affiliations:** Department of Bioengineering, University of Illinois Urbana-Champaign, Urbana, IL 61801, USA; Center for Artificial Intelligence and Modeling, Carl R. Woese Institute for Genomic Biology, University of Illinois Urbana-Champaign, Urbana, IL 61801, USA; Department of Physics, University of Illinois Urbana-Champaign, Urbana, IL 61801, USA

## Abstract

Many microbial communities in nature are complex, with hundreds of coexisting strains and the resources they consume. We currently lack the ability to assemble and manipulate such communities in a predictable manner in the lab. Here, we take a first step in this direction by introducing and studying a simplified consumer resource model of such complex communities in serial dilution experiments. The main assumption of our model is that during the growth phase of the cycle, strains share resources and produce metabolic byproducts in proportion to their average abundances and strain-specific consumption/production fluxes. We fit the model to describe serial dilution experiments in hCom2, a defined synthetic human gut microbiome with a steady-state diversity of 63 species growing on a rich media, using consumption and production fluxes inferred from metabolomics experiments. The model predicts serial dilution dynamics reasonably well, with a correlation coefficient between predicted and observed strain abundances as high as 0.8. We applied our model to: (i) calculate steady-state abundances of leave-one-out communities and use these results to infer the interaction network between strains; (ii) explore direct and indirect interactions between strains and resources by increasing concentrations of individual resources and monitoring changes in strain abundances; (iii) construct a resource supplementation protocol to maximally equalize steady-state strain abundances.

## INTRODUCTION

Many microbial communities in both natural^1–8^ (human gut, plant rhizosphere, soil, ocean) and artificial or synthetic^9–14^ settings are highly diverse, with the number of coexisting strains reaching into the hundreds. Such diversity of strains is often accompanied by the diversity of nutrients, with several hundred resources necessary to support the growth of hundreds of strains. In addition, many natural environments and most lab experiments on microbial communities are characterized by “boom-and-bust” cycles, where nutrients are added in large batches at the beginning of each cycle and species either die or get diluted in large ratios at the end of the cycle. Such serial dilution experiments are very easy to perform in the lab, but relatively difficult to model predictively^15–18^

There are a number of approaches suitable for modeling microbial communities with different levels of diversity. One of the most popular mathematical techniques uses generalized Lotka-Volterra (gLV) models^19–21^, where strains are assumed to directly interact with each other. In gLV models, the exponential growth rate for each strain is approximated as a linear combination of the abundances of all strains in the community. However, this strategy does not explicitly account for resource competition or metabolic cross-feeding between strains. Thus, gLV models have been shown to be inadequate for the modeling of microbial communities with such indirect interactions^22^.

Consumer-Resource Models (CRMs), such as the MacArthur model^23–25^ or the Tilman model^26,27^ are another popular choice for modeling microbial communities. These models explicitly describe shifts in community composition in response to changes in resource supply rates. CRMs also explicitly account for the consumption and production of metabolites by species. However, species and resources in the steady state of a classical CRM are assumed to be constant in time, which is appropriate for chemostat-like stable environments, but not for boom-and-bust dynamics.

The new generation of CRMs was developed to model microbial community dynamics in serial dilution and other strongly fluctuating environments^15–18,28^. CRMs with different levels of resolution are appropriate to describe communities with different levels of complexity. For example, we and others have developed the most detailed CRMs that can be used to describe low complexity communities with fewer than six strains and resources^15–17,29,30^. These models explicitly account for a variation in depletion times of individual resources and differences in growth rates and time lags of strains in each of the resulting temporal niches.

At the intermediate level of community complexity, Ho and collaborators recently proposed a CRM^28^ for a simplified synthetic human gut microbiome consisting of 15 representative strains. Even at this reduced level of diversity, a number of approximations and simplifications were necessary. The authors clustered resources into 2^15^ groups based on the exact subset of species capable of consuming them, and then selected about 30 of the most abundant binary groups. However, this method is not scalable to the most complex communities with hundreds of species such as, e.g., hCom2 - a synthetic human gut microbiome with 120 strains developed in Ref.^11^.

Here, we present a simplified mechanistic model of complex microbial communities with hundreds of coexisting strains and resources. To test the performance of our model, we use it to predict the dynamics of 63 strains from hCom2 surviving in a serial dilution experiment^31^. The hCom2 community has been developed as a potential candidate for gut microbiome transplantation therapy^11^. It is therefore of practical importance to develop a reliable predictive model that can be used, for example, to predict the response of this synthetic community to various perturbations, or to attempt to equalize strain abundances prior to transplantation into the patient^32^.

After fitting the parameters of the model based on the metabolomics data of Ref.^33^ and serial dilution data from Ref.^31^, we carried out three in silico perturbation experiments. First, we performed a leave-one-out experiment in which individual strains were removed from the inoculum one by one. This allowed us to resolve strain-strain interactions in the community as either cooperative or competitive. In the second experiment, we perturbed individual metabolites by increasing their concentrations in the medium and showed that this could both increase and decrease strain abundances. While an increase in the abundance of a strain in response to an increase in the concentration of a metabolite in the medium can be caused by both direct consumption of that metabolite and indirect effects such as cross-feeding on metabolic byproducts of other strains, a decrease in the abundance of a strain can only be caused by indirect effects such as competition for that and other metabolites with the rest of the strains in the community. Finally, based on the results of the second experiment, in the third experiment we proposed and implemented a greedy algorithm aimed at equalizing strain abundances in the community by increasing the concentrations of multiple metabolites.

## MODEL AND RESULTS

### Consumer resource model of a complex community in serial dilution experiments

We introduce a simplified Consumer Resource Model (CRM) to predict the dynamics of a complex community composed of multiple microbial strains in the presence of multiple resources in serial dilution experiments. Consider *n*_*S*_ strains growing on *n*_*R*_ resources. The strains are grown in a serial dilution experiment, where the community culture is serially passaged. At the beginning of a passage, all strains are diluted by the same factor *D*. After that, the strains grow exponentially while consuming different resources. The abundance of a resource *i* is given by *R*_*i*_, measured in units of its contribution to biomass. Thus we assume that yields of the same resource to the biomass of different strains are equal to each other and without loss of generality can be rescaled to 1. While the resource *i* is present in the environment, it is consumed by the strain *α* in proportion to its abundance *N*_*α*_(*t*) and strain-specific consumption flux, *c*_*α-i*_ calculated per unit of its biomass. When multiple strains consume the same resource, the fraction consumed by the strain *α* is given by 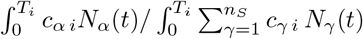. Here *T*_*i*_ is the time since the start of the growth cycle when the resource *i* is depleted. With these assumptions, mass conservation at the end of the growth cycle can be written as

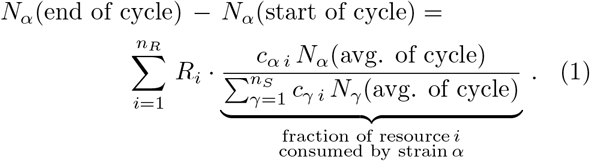

Above we assumed that at any passage of a serial dilution experiment all resources get completely depleted. *N*_*α*_(start of cycle) and *N*_*α*_(end of cycle) are the abundances of strain *α* at the start and end of the growth cycle, respectively. *N*_*α*_(avg. of cycle) is the average strain abundance during the growth cycle, which determines the proportions for sharing resources among the strains. In principle, resources are consumed throughout the growth cycle, and an accurate calculation of *N*_*α*_(avg. of cycle) requires integration of the instantaneous abundance over the entire growth cycle, but these data are usually not available. Assuming exponential growth with an approximately constant growth rate between 0 and *T*_*i*_, we approximate *N*_*α*_(avg. of cycle) by the geometric mean of *N*_*α*_(start of cycle) and *N*_*α*_(end of cycle):

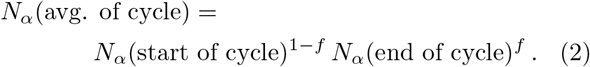

Here a single parameter, *f ∈* [0, 1], which we call the time fraction, appoximately accounts for two effects; (i) the fact that some nutrients get depleted prior to the end of the growth cycle, when the last nutrient gets depleted, (ii) the fact that *N*_*α*_(start of cycle) *< N*_*α*_(avg. of cycle) *< N*_*α*_(end of cycle). For exponentially growing species, all resources are depleted towards the end of the growth cycle, thus we expect *f* to be close to 1. In principle, depletion times *T*_*i*_ differ between resources, so the time fraction *f*_*i*_ should also be resource specific. However, when we tested our model on serial dilution data in a complex synthetic human gut community fitting resource-specific values of *f*_*i*_ resulted in overfitting. Therefore, we simplified the model to use the same time fraction *f* for all resources. Later, we demonstrate a simple method for estimating *f* from the experimental data.

At the beginning of a passage, after the end of the previous passage, all strain abundances are diluted by the factor *D*. Then, for any two consecutive passages, *k −* 1 and *k, N*_*γ*_(start of cycle *k*) = *N*_*γ*_(end of cycle *k −* 1)*/D*. Now, substituting this and Eq. (2) into Eq. (1), we get Eq.(3). There, we have also introduced interactions between strains through cross-feeding, and this is implemented as a multiplicative factor converting the concentration *R*_*i*_ of a resource *i* in the bolus medium to its total concentration ultimately consumed by the strains *R*^(total)^. The multiplier described by Eq. (3b) depends on the normalized production flux, *p*_*γ-i*_, calculated per unit of biomass of the strain *γ* producing the resource *i*. If a resource *i* is not produced as a metabolic byproduct by any strain, then *p*_*γ-i*_ = 0, and *R*^(total)^ = *R*_*i*_. However, if *i* is produced by at least one strain, then *p*_*γ-i*_ *>* 0 for some *γ*, and *R*^(total)^ *> R*_*i*_. If *D ≫* 1, then the second term on the left hand side (LHS) in Eq. (3a) can be omitted. Eq. (3) is the version of the model used for all analyses in the paper.

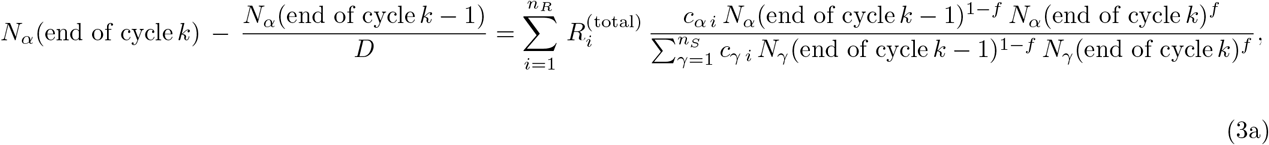

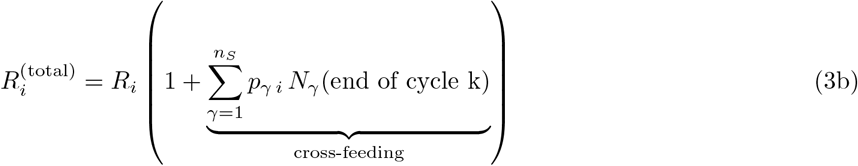

Our treatment of cross-feeding interactions is again a necessary simplification of what is generally a more complex dynamic process. We ignore possible differences in the timing of the production of metabolic by-products by using the biomass of the producing strains at the end of the cycle as a multiplier in Eq. (3b). This approximation is justified in the case of rapid exponential growth where the average biomass is close to its value at the end of the cycle. Solving Eqs. (3a) and (3b) for *N*_*α*_(end of cycle *k*) as a function of *N*_*α*_(end of cycle *k −* 1) defined passage-to-passage dynamics in serial dilution experiments. One can also set *N*_*α*_(end of cycle *k*) = *N*_*α*_(end of cycle *k −* 1) and solve for steady-state abundances reached after multiple passages.

An illustration of the model is shown in Fig. 1. In this example, there are two strains, *A* and *B*, and three resources, *R*_1_, *R*_2_, and *R*_3_. Both strains consume *R*_1_ and produce *R*_2_ (Fig. 1a). However, neither strain consumes or produces *R*_3_ (Fig. 1a). *R*_1_ is divided among the strains in proportion to their average abundances, *N*_*A*_(avg.) and *N*_*B*_(avg.), respectively, weighted by the respective consumption fluxes for *R*_1_, *c*_*A* 1_ and *c*_*B* 1_ (Fig. 1b). The cross-feeding term involving *R*_2_ is proportional to the abundances of strains *A* and *B* at the end of the growth cycle, *N*_*A*_(end) and *N*_*B*_(end), respectively. The contributions of *A* and *B* to the cross-feeding term are also weighted by the respective production fluxes for *R*_2_, *p*_*A* 2_ and *p*_*B* 2_ (Fig. 1c).

**FIG. 1.**
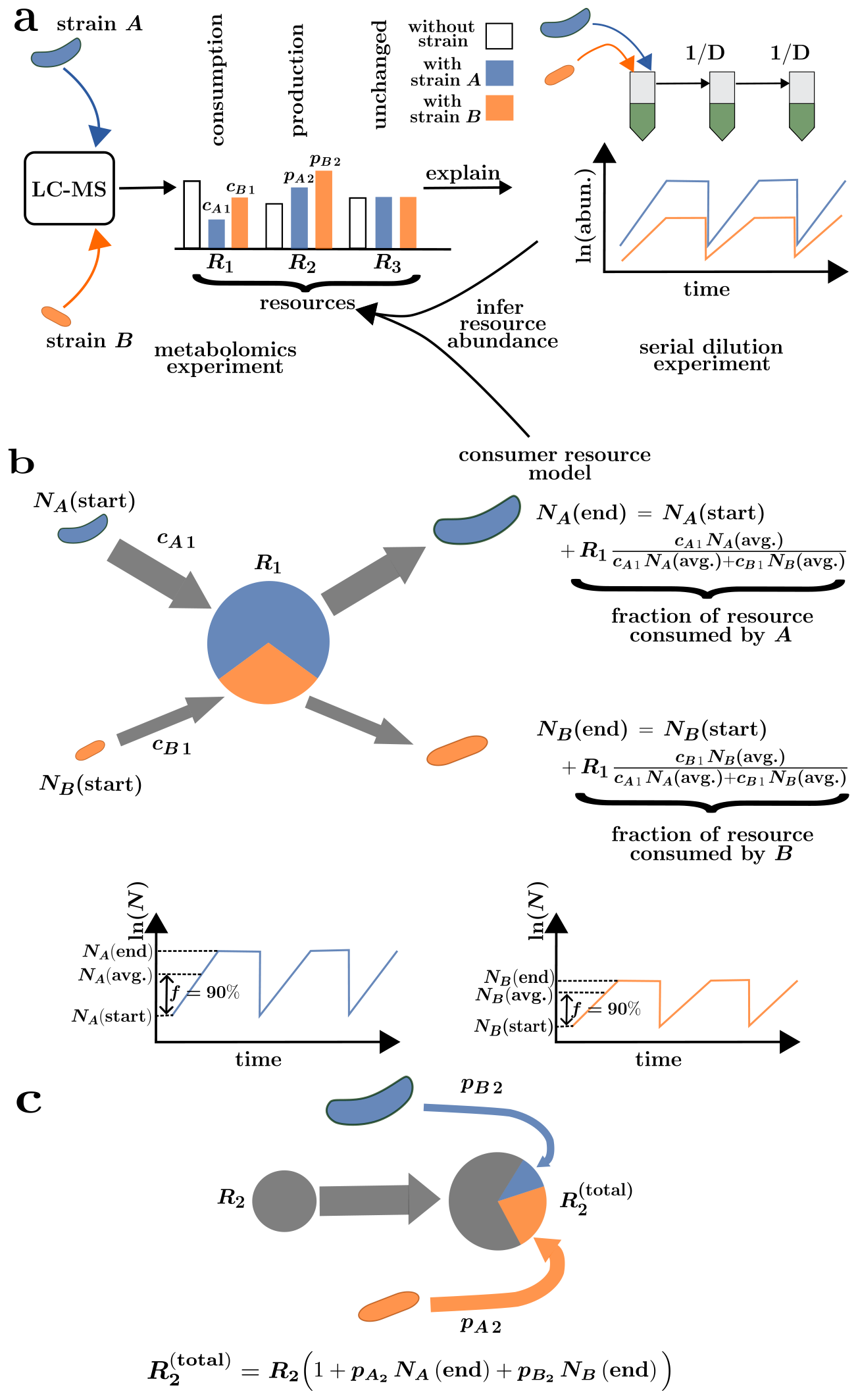
A schematic of our consumer resource model. This example has 2 strains (*A* and *B*) and 3 resources (*R*_1_, *R*_2_, and *R*_3_) in a hypothetical community. **a)** (Left) Production and consumption fluxes are measured on a strain-by-strain basis from, e.g. a metabolomics experiment using a liquid chromatography-mass spectrometry (LC-MS) technique. A unit of biomass of strains *A* and *B* consumes *R*_1_ with consumption fluxes *c*_*A* 1_ and *c*_*B* 1_, respectively, and produces *R*_2_ as a byproduct with production fluxes *p*_*A* 2_ and *p*_*B* 2_, respectively. Neither strain consumes or produces *R*_3_. (Right) Abundances for the strains in the community at multiple passages are obtained from a serial dilution experiment where abundances are diluted by a factor *D* at the beginning of the growth cycle. **b)** According to the model, during the growth cycle for each passage of the serial dilution experiment, *R*_1_ is divided between *A* and *B* in proportion to their average abundances *N*_*A*_(avg.) and *N*_*B*_ (avg.), respectively, and to their consumption fluxes *c*_*A* 1_ and *c*_*B* 1_, respectively. The average strain abundances are obtained as the geometric mean of the abundances at the beginning and end of the growth cycle as given by Eq. (2). **c)** According to the model, for all strains in the community that consume *R*_2_, its excess concentration is contributed by its production as a byproduct by *A* and *B*. The contributions of *A* and *B* to this excess are proportional to their abundances at the end of the growth cycle *N*_*A*_(end) and *N*_*B*_ (end), respectively, and to their normalized production fluxes *p*_*A* 2_ and *p*_*B* 2_, respectively.

### Application of the model to predict serial dilution dynamics of a complex synthetic human gut microbial community

#### Data for fitting the consumer resource model

We have applied our model in Eq. (3) to explain the serial dilution dynamics of a complex synthetic human gut microbial community hCom2^31^ grown in a rich media (mega media) using two published and publicly available data sets^31,33^. Both datasets contain strains of a defined hCom2 community that was initially populated with prevalent bacterial strains from the human gut microbiome and subsequently challenged with a human fecal sample to fill open niches, resulting in increased stability to fecal challenge and robust colonization resistance (see Ref.^11^ for details). One of the potential applications of such a community is gut microbiome transplantation or supplementation, which may have therapeutic implications for various diseases.

Fitting our model to hCom2 involved estimating resource abundances *R*_*i*_, which in this case represent different metabolites. To infer *R*_*i*_, we collected consumption and production fluxes (*c*_*α-i*_ and *p*_*γ-i*_, respectively) of 63 strains for 292 metabolites from the metabolomics experiments of Han et al. 2021^33^. The names of the 63 strains along with the abbreviations we assigned are given in table S1. Details on the data set and the calculation of *c*_*α-i*_ and *p*_*γ-i*_ are given in the Methods section.

In addition, we obtained the strain abundances *N*_*α*_(end of cycle) for hCom2 grown in mega media for multiple passages of a serial dilution experiment from Jin et al. 2023^31^. For model fitting, we used changes in strain abundances between the first and the second passage of the serial dilution experiment. This passage had the most dynamics of all the passages where we could separately study biological replicates^31^ and was therefore the most suitable for fitting our dynamic model. The change from inoculum to first passage was less suitable because the inoculum data had only two biological replicates and these were not matched to three biological replicates in the first passage. We used 189 data points (three biological replicates for the abundances of 63 strains) to fit Eq. 3 for *R*_*i*_. Details for this dataset are also given in the Methods section.

#### Coarse-graining the metabolite consumption and production fluxes

As discussed above, the number of available data points to fit the model is 189, which is less than the number of metabolites, 292. To avoid overfitting, we coarse-grained the metabolites by clustering them, which resulted in 98 clusters, including 10 non-singleton and 88 singleton clusters. The average consumption and production fluxes of the 63 strains were then calculated for these 98 metabolite clusters. The clustering process and the estimation of consumption and production fluxes for the metabolite clusters are described in detail in the Methods section. A hierarchically clustered heatmap for the 98 metabolite clusters is shown in Fig. S2. The names of the metabolites in each metabolite cluster are given in Table. S2. In the rest of the manuscript, we use the terms metabolite and metabolite cluster synonymously, unless otherwise indicated.

#### Estimating the time fraction f

For the synthetic human gut microbial community hCom2, we performed a systematic search for the time fraction *f* used in Eqs. (2) and (3) and found that the predictive performance of our fitted model shows a sharp peak around *f* = 0.9 (Methods, Fig. S1). This is expected for rapidly growing bacteria and experiments with large dilution factors (*D* = 15, 000 in Ref.^31^), where resources tend to be depleted towards the end of the growth cycle. Details of the estimation procedure are given in the Methods section.

#### Fitting and validating the model on the synthetic human gut microbiome

After obtaining the consumption and production fluxes, time fraction, and using abundances at passages 1 and 2 of the serial dilution experiment, we inferred *R*_*i*_ for the metabolite clusters by plugging these values into our model in Eq. (3). Details of the fitting procedure are given in the Methods section. Of the 98 metabolite clusters, 38 had non-zero (*≥* 10^*−*8^) fitted *R*_*i*_.

Using the fitted values of *R*_*i*_, we validated the model through an independent in silico prediction experiment, which consisted of predicting the dynamics of strain abundances all the way from the inoculum to steady state reached around passage 3. To do this, we started with the experimentally measured strain abundances in the inoculum (average over two biological replicates in Ref.^31^). Since the inoculum data were not used in our fitting procedure, starting from it helps to obtain an independent estimate of model performance. Using the inoculum abundances as *N*_*α*_(end of cycle 0) Eq. (3) with with the inferred *R*_*i*_ was iteratively solved to obtain *N*_*α*_(end of cycle 1), *N*_*α*_(end of cycle 2), and *N*_*α*_(end of cycle 3). The details of solving Eq. (3) for prediction are described in the Methods section. We stopped the model at passage 3, since the experimental data suggest that the community reaches steady state by then^31^.

A comparison between the predicted and observed abundances from the above in silico experiment is shown for passages 1 to 3 in Fig. 2. The predicted and observed abundances are strongly correlated at passages 2 (Pearson’s correlation = 0.77, *p* = 1.58 *×* 10^*−*13^) and 3 (Pearson’s correlation = 0.77, *p* = 1.34 *×* 10^*−*13^). The correlation at passage 1 is somewhat lower (Pearson’s correlation = 0.58, *p* = 8.24*×*10^*−*7^). This suggests that the model is better at predicting abundances at later passages, including steady state, than at the first passage.

**FIG. 2.**
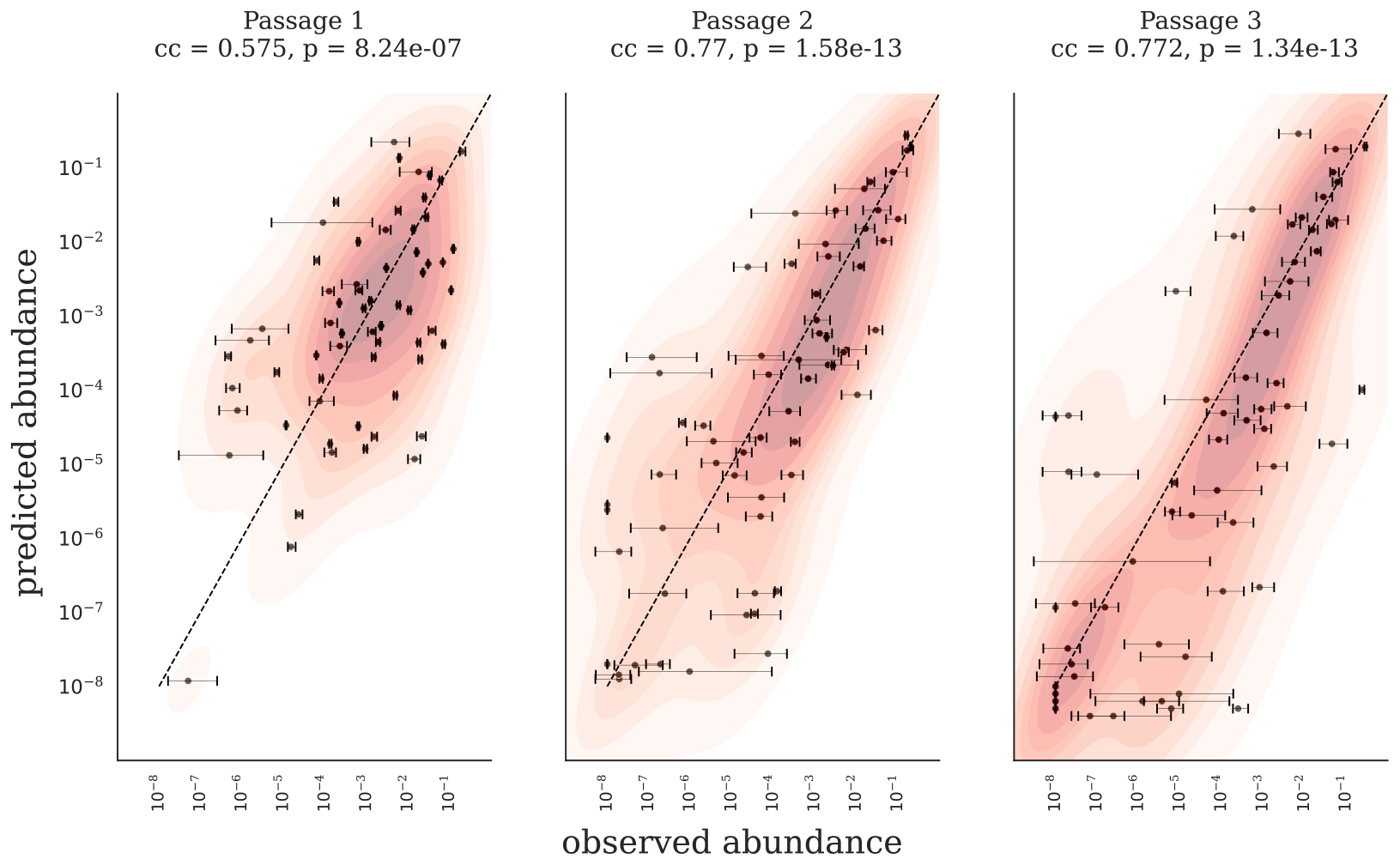
The model predicts serial dilution dynamics for a complex synthetic human gut microbiome. The model for hCom2 was validated by predicting strain abundances at different passages from inoculum abundances. Pearson’s correlation coefficients (cc) and p-values between log_10_ of predicted and observed abundances are listed above each panel. Each point on the scatterplot represents one strain. Error bars correspond to the range (maximum minus minimum) of observed strain abundances across three biological replicates.

#### Accounting for variability in biological replicates

In addition to the correlation coefficient, Root Mean Squared Error (RMSE), on the log_10_ scale, is another metric that can be used to quantify model performance. RMSE measures how much, on average, the predicted strain abundances deviate from the observed strain abundances on the log scale, and unlike correlation, a lower value for RMSE is desirable. We found that the RMSE for the first passage was lower than for the other two (Fig. S3), suggesting that the model performance was better for the first passage. This is contrary to the trend we saw for correlation, where performance was lower for the first passage. We hypothesized that this was a consequence of the relatively low replicate-to-replicate variability in the observed data at the first passage. To quantitatively test this hypothesis, we estimated the biological replicate-to-replicate variability by calculating the RMSE between biological replicates at each passage. We found that a significant portion of the residual error in our predictions was due to variability between biological replicates (Fig. S3). For passages 2 and 3, the biological replicate-to-replicate variability accounted for more than 50% of the RMSE for our model predictions (Fig. S3). Now, the adjusted RMSE (RMSE minus biological replicate-to-replicate variability) was higher for the first passage compared to the other two (Fig. S3). Thus, the prediction performance of our model was better at later passages, which is consistent with the trend from the correlation analysis.

Such large biological replicate-to-replicate variability can result from undefined media, such as the mega media used in Ref.^31^ with significant variation in resource concentrations in the media between replicates and/or passages. Furthermore, in Ref.^33^, the same mega-media was used to measure consumption and production fluxes, resulting in significant day-to-day variation in *c*_*α-i*_ and *p*_*γ-i*_ measurements. This can contribute to the residual error in the model predictions.

The width of the error bars in Fig. 2 appears to be larger for the low abundance strains compared to the high and medium abundance strains. We tested this quantitatively and found that the width of the error bars is indeed negatively correlated with strain abundances (Fig. S4d-f). Another measure of the same effect is the RMSE between biological replicates, which also decreased steadily with increasing abundance (red curves in Fig. S4a-c). Here, biological replicate-to-replicate variability was calculated as the cumulative RMSE between biological replicates over a subset of strains. We subset the strains by thresholding the average observed abundances at steady state (details are given in the Methods section). Based on this observation, we hypothesized that the error in model predictions should also improve with increasing abundance. To test this, we computed the cumulative RMSE for the model predictions by thresholding the average observed abundances at steady state (details are given in the Methods section).. For all passages, we found that the RMSE between predicted and observed abundances decreased steadily as we moved from low to high abundance strains (blue curves in Fig. S4a-c). The residual error of our predictions (the difference between the blue and red curves in Fig. S4a-c) also tended to become smaller for high-abundance strains that had low biological replicate-to-replicate variability.

### In-Silico Experiments on the Synthetic Human Gut Microbial Community

After validating our model, we used it to perform three in silico experiments to study the response of the synthetic gut community to different perturbations: In (i), we removed one strain from the inoculum and predicted the steady-state abundances of the remaining strains. By repeating this leave-one-out experiment for each of the 63 strains, we calculated the network of significant direct and indirect interactions between strains; In (ii), we increased the concentrations of individual resources 100-fold and examined the resulting changes in steady-state strain abundances. This experiment produced a matrix of direct and indirect interactions between resources and strains. Encouraged by experiments (i) and (ii), we tested our ability to manipulate the community through multi-nutrient supplementation. In (iii), we increased the concentrations of several resources to balance the abundances of strains as much as possible. The details of these three in silico experiments are described in the following sections.

#### Sensitivity of steady-state strain abundances to removal of a single strain from the inoculum

The first perturbation experiment we performed was to remove one strain from the inoculum and predict the steady-state abundances of the remaining strains. Each of 63 strains was removed in a different leave-one-out experiment. We compared the predicted steady-state abundances between the perturbed and unperturbed communities. This allowed us to estimate interactions between strains in the community, and the robustness of the community to extinction-driven perturbations. We found two types of interactions between strains - 1) cooperative (positive), and 2) competitive (negative). We represent these interactions as a directed graph in Fig. 3a. For each edge, the source and destination nodes represent the removed and perturbed strains, respectively. To filter out noise, we kept only those edges where the abundance of the perturbed strain either increased more than 10-fold (red arrows in Fig. 3a) or decreased more than 10-fold (green arrows in Fig. 3a). Each edge is weighted by the absolute amount of change in abundance (on the log_10_ scale) of the perturbed strain when the strain at the source of the edge is removed. Removing 11 of the 63 strains in independent leave-one-out experiments resulted in a significant shift in the abundance of at least one perturbed strain. Of the 63 strains, 35 were perturbed by the removal of at least one strain. Node sizes were set proportional to the weighted in-degree (sum of weights of all incoming edges at a node).

**FIG. 3.**
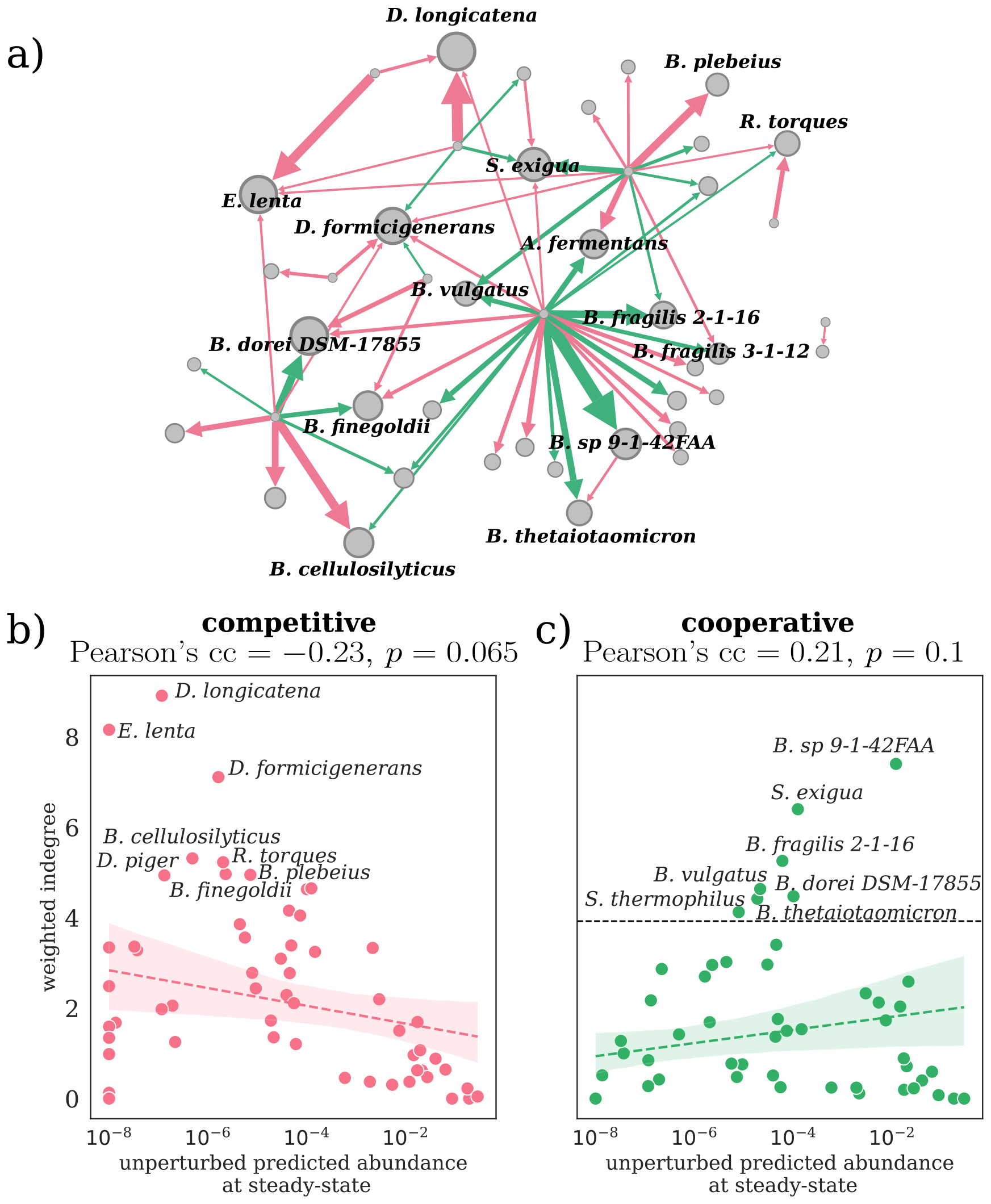
Leave-one-out perturbations reveal competitive and cooperative interactions between strains in the synthetic gut community. **a)** The directed strain-strain interaction network from leave-one-out experiments displays interaction edges directed from the removed strain to the perturbed strain and weighted by the log10-scaled abundance change of perturbed strains. Node sizes reflect their total weighted in-degrees, with the top 15 strains labeled. Cooperative and competitive interactions are depicted as green and red edges, respectively. **b)** Weighted in-degrees of strains from competitive edges (red) plotted against their steady-state abundances in the unperturbed community. The top 8 strains most sensitive to competitive interactions are labelled. **c)** Similar to **b)**, but with in-degrees for cooperative edges (green), with labels for the top 7 strains most sensitive to cooperative interactions. Panels **b)** and **c)** include Pearson’s correlation coefficients between frequency and weighted degree, along with a dashed line fit and a shaded confidence interval, none of which are significant at the 95% level.

For cooperative interactions (green edges in Fig. 3a), removal of a strain leads to a decrease in the abundance of the perturbed strain. One way this can happen is if the strain with deceased abundance cross-feeds on the metabolites produced by the removed strain. Consequently, when the strain is removed, the perturbed strain has fewer resources and responds with a decrease in abundance. To test this hypothesis, we calculated a cross-feeding score by counting the fraction of metabolites consumed by the perturbed strain that were produced by the removed strain (see Methods section for details on calculating the cross-feeding score). The score is equal to 1 when all metabolites produced by the removed strain are consumed by the perturbed strain and 0 when none of them are consumed. For cooperative or green edges, we found that, the average cross-feeding score of of 0.64 was significantly higher (*p* = 9.7 *×* 10^*−*5^, two-tailed Mann-Whitney Wilcoxon test) than the average cross-feeding score of 0.55 between pairs of non-interacting strains or for strains connected by competitive interactions (red edges in Fig. S7.

Next, we analyzed the competitive edges in the interaction network. In a competitive interaction, the removal of one strain leads to an increase in the abundance of another strain. In our model, the primary source of these interactions is competition for resources between the removed and perturbed strains. We calculated a competition score by counting the fraction of resources (or their clusters) consumed by the perturbed strain that were also consumed by the removed strain (see Methods section for details on calculating the competition score). For competitive edges, the average competition score of 0.65 between the removed and perturbed strains was significantly larger (*p* = 0.02, two-tailed Mann-Whitney Wilcoxon test) than the average competition score of 0.58 between non-interacting pairs or cooperative interactions (green edges) in Fig. S7.

The strain-strain interaction network’s properties were analyzed using weighted out- and in-degree metrics from network analysis. We observed that strains with high outdegree, indicating a substantial impact on the community when removed, often correlated with high abundance in the unperturbed community (Fig. S5). This pattern persisted across both cooperative and competitive interactions (Fig. S5a, b). The removal of a dominant strain typically results in resource reallocation, allowing competitors to expand and alter the community’s relative abundance. Notably, this community shift is intrinsic and not an artifact of the normalization process in relative abundance studies, as demonstrated in Fig. S6a, where the exclusion of the most abundant strain affects other strains by different amounts.

We examined the weighted in-degree distribution in the strain-strain interaction network, considering both cooperative and competitive edges. The total weighted in-degree of a strain represents its overall sensitivity to the removal of other strains. In Fig. 3a, node sizes are scaled to their total weighted in-degree. Figs. 3b, c display the average weighted in-degree, segregated by edge type, against the unperturbed abundance of strains at steady state. For competitive edges, there’s a negative correlation between average in-degree and the log_10_ of strain abundance (Pearson’s cc of *−*0.27, *p* = 0.03, Fig. 3a). This trend aligns with the resource conservation law, where the removal of a low abundance strain insignificantly impacts higher abundance strains due to limited resource reallocation. The greater the abundance of the strain, the smaller the number of other strains that can potentially affect it, and thus the lower the expected in-degree,

Medium abundance strains showed a higher tendency for cooperative interactions (Fig. 3c). Notably, 5 out of 7 of these strains were from the genus *Bacteroides* (labeled in Fig. 3a, c), representing a significant 71.4% occurrence compared to their 33.3% fraction in the total set of 63 strains (*p* = 0.036, one-sided hypergeometric test). This trend persists even at a lower cutoff, with *Bacteroides* comprising 9 of the top 15 most responsive strains (*p* = 0.0154, one-sided hypergeometric test). However, this pattern is limited to *Bacteroides* strains with intermediate abundances. In fact, the top 10 most abundant strains, including three *Bacteroides* species (*Bacteroides uniformis* ATCC-15579, *Bacteroides thetaiotaomicron* VPI-5482, and *Bacteroides stercoris* ATCC-43183) have exactly zero weighted in-degree.

#### Sensitivity of the steady-state strain abundance to increase in the concentration of a single metabolite

Leave-one-out experiments show competition and cooperation between strains without identifying the responsible metabolites. To further understand this, we analyzed how the steady-state abundance of each strain changes when the concentration of a single metabolite is increased. Since decreasing metabolite concentrations in undefined mega media is impossible in practice, we focused on sensitivity to increases. For each experiment, we increased the concentration of one of 38 metabolite clusters by a factor of 100. Then, using Eq. (3), we predicted the new steady-state strain abundances. The logarithmic scale (base 10) difference between the perturbed and unperturbed abundances for each strain indicated its sensitivity to that metabolite.

Fig. 4 presents a clustered heatmap illustrating how strain abundances respond to metabolite concentration perturbations. These perturbations caused both increases (33%) and decreases (67%) in strain abundances, revealing distinct clusters. For instance, strains *Coprococcus comes* ATCC 27758, *Holdemanella biformis* DSM 3989, *Dorea longicatena* DSM 13814, and *Anaerotruncus colihominis* DSM 17241 showed notable abundance increases in response to several metabolites, possibly due to direct consumption or indirect effects like cross-feeding. Conversely, a cluster of mainly *Bacteroides* strains at the heatmap’s top exhibited abundance decreases in response to 10 specific metabolites, likely from competition-related indirect effects. Some strains remained largely unaffected by metabolite perturbations. For example, the following 10 strains marked with a black arrow in Fig.4 were perturbed by fewer than 3 metabolites: *Bacteroides caccae* DSM 43185, *Marvinbryantia formatexigens* DSM 14469, *Bacteroides coprophilus* DSM 18228, *Clostridium leptum* DSM 753, *Ruminococcus bromii* ATCC 8503, *Parabacteroides johnsonii* DSM 18315, *Lactococcus lactis* DSMZ 20729, *Lactobacillus ruminis* ATCC 25644, *Tyzzerella nexilis* DSM 2243, and *Catenibacterium mitsuokai* DSM 15897. Consequently, we excluded these strains from our subsequent in silico experiment, which focuses on equalizing steady-state strain abundances through increasing concentrations of multiple metabolites (detailed in the following section).

**FIG. 4.**
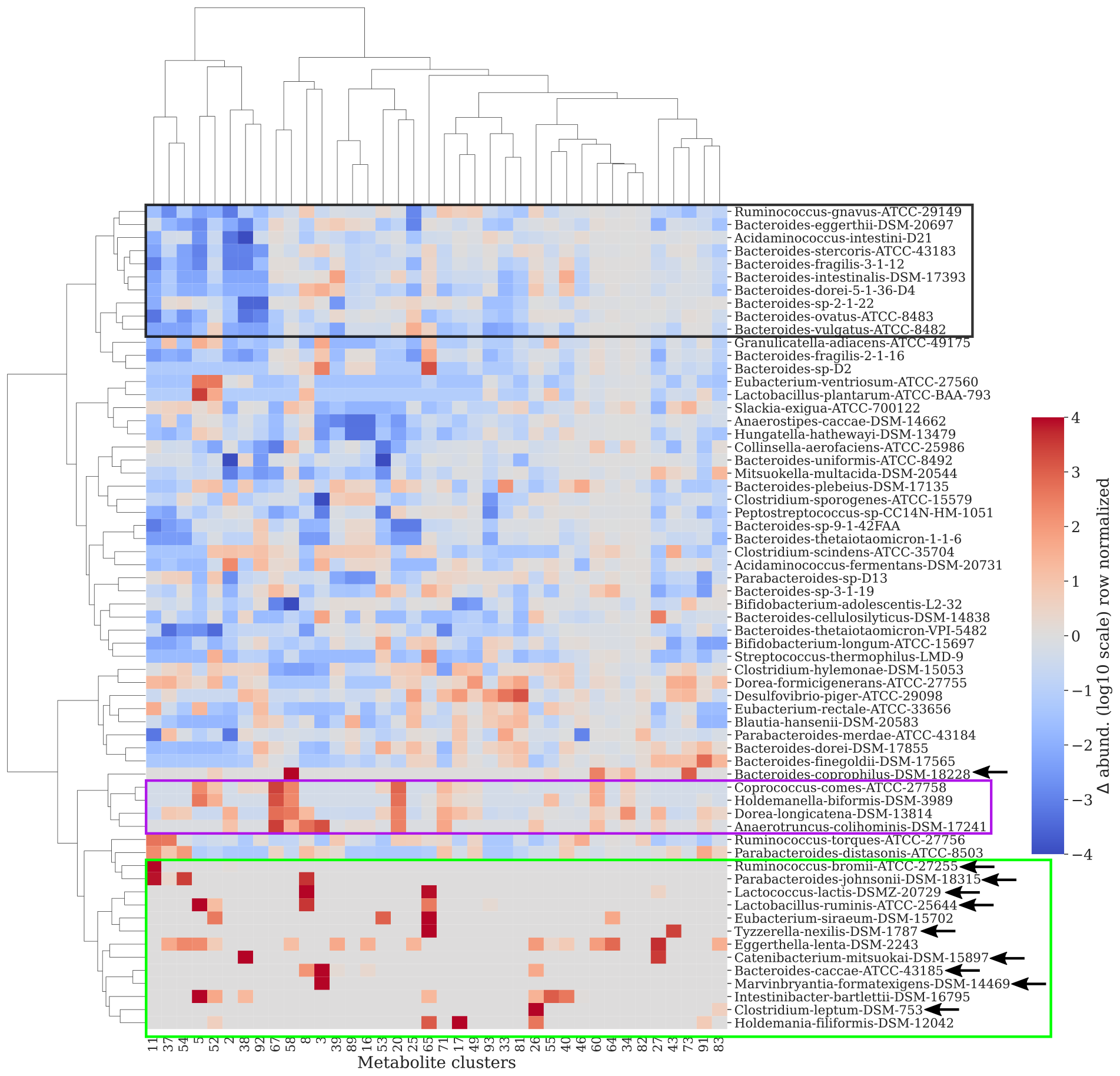
Response of strain abundances to increases in concentrations of individual metabolites. A hierarchically clustered heatmap of the ratio (on log_10_ scale) between perturbed and unperturbed strain abundances at steady state in response to an increase in the concentration of a single metabolite in the mega media. Columns and rows represent 38 metabolites (or their groups) and 63 strains in the synthetic gut community, respectively. Three representative clusters of strains have been enclosed in boxes for easy identification. 10 strains marked with a black arrow were classified as poorly responsive to metabolite addition and were therefore excluded from the in silico experiment to equalize the strain abundances in Fig. 5.

#### Supplementing the mega media with multiple metabolites can approximately equalize strain abundances

In the previous in silico experiment we saw that perturbations in individual metabolites can both increase and decrease strain abundances in hCom2. This suggests that simultaneous perturbations in multiple metabolites may be able to, at least approximately, equalize strain abundances. Equalizing abundances has practical implications for gut microbiome transplantation therapy. If abundances can be equalized, then all strains in the transplanted community would have an equal chance to survive and colonize the gut.

To accomplish this we designed a greedy algorithm in which we changed abundances of multiple metabolites one-by-one. At every step we changed the abundance of a single metabolite that brings the community closest to uniformity. For this experiment we removed the 10 non-responsive strains identified in the previous section and marked with a black arrow in Fig. 4. Details of the greedy algorithm are given in the Methods section. Since the greedy algorithm was stochastic, we repeated it 10 times to obtain 10 possible metabolite perturbation profiles and 10 different perturbed *R*_*i*_.

We identified 56 metabolites that were altered in at least one of the 10 runs of the greedy algorithm. The abundances for these 56 metabolites are shown in a clustered heatmap in Fig. S8. 20 of the 38 non-zero unperturbed *R*_*i*_ are present in this list. The remaining metabolites had *R*_*i*_ = 0 before perturbation. We averaged the perturbed *R*_*i*_ over the 10 runs of the greedy algorithm and used this average to calculate the steady-state abundances starting from the inoculum for hCom2. The community is now much closer to uniformity compared to the unperturbed scenario (Fig. 5). The log_10_-RMSE between the perturbed steady state and perfect equalization was 1.7, which was two times smaller than the log_10_-RMSE of 3.4 between the unperturbed steady state and perfect equalisation.

**FIG. 5.**
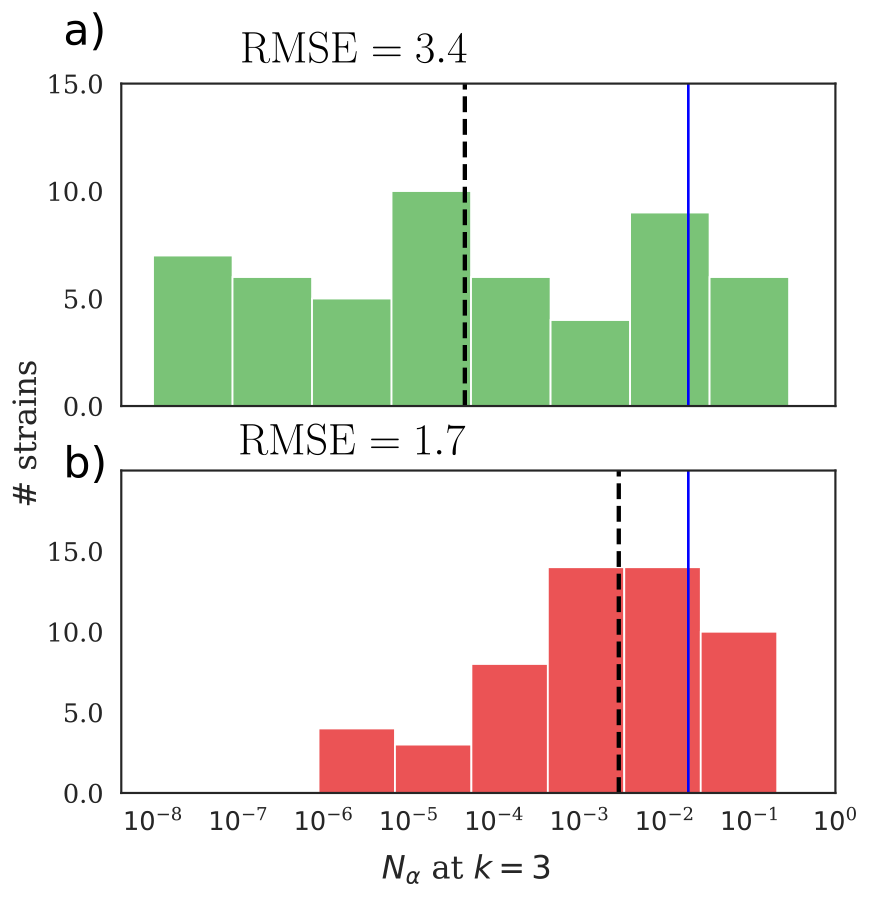
Equalisation of steady-state strain abundances for hCom2. Distribution of steady-state strain abundances for hCom2 without (**a)**) and with (**b)**) perturbation aimed at equalizing strain abundances. As a result of this perturbation, the *log*_10_-RMSE of the steady state abundances decreased twofold from 3.4 to 1.7. The blue solid lines show the ideal steady state abundance, the dashed black lines show the mean abundance values for the unperturbed and equalized communities.

## DISCUSSION

Here, we introduced and studied a simplified consumer resource model for complex microbial communities with hundreds of coexisting strains growing on several hundred resources in serial dilution lab experiments. The central assumption of the model is that during the growth phase of the cycle, strains share resources in proportion to their average abundances and strain- and resource-specific consumption fluxes. This assumption was applied to all resources in the rich media to link strain abundances at successive passages of serial dilution experiments via mass conservation. We also incorporated cross-feeding into the model via a simple linear term linking the abundances of strains producing a given resource to its excess concentration in the medium. Consumption and production fluxes can be inferred from metabolomics performed on batch growth experiments with individual strains. Our model can then be fitted to the data in a serial dilution experiment to infer resource concentrations, *R*_*i*_ *≥* 0, by solving Eq. (3) for a single dilution cycle as a non-negative least squares (NNLS) problem. With the fitted *R*_*i*_, the model can be used to predict the dynamics at other dilution cycles not used in the initial fit.

We tested the model on a defined synthetic human gut microbiome, hCom2^11^ growing on a rich medium with several hundred metabolites reaching steady-state diversity of around 60 strains^31^. To fit the model to hCom2, we first obtained strain-specific resource consumption and production fluxes in the mega medium from the metabolomics experiments described in Han et al. 2021^33^. The abundances of strains in hCom2 grown in mega medium at multiple passages of serial dilution experiments were obtained from Ref.^31^.

Modeling a complex synthetic community such as hCom2 requires several simplifying assumptions. First, our model assumes that multiple strains consuming the same resource share it in proportion to their average abundances during the growth cycle. An accurate calculation of the average abundance requires integration of the instantaneous abundance over the entire growth cycle, but these data were not available in Ref.^31^. Since strains grow approximately exponentially, we approximated the average abundance by the geometric mean of the strain abundances at the beginning and end of the growth cycle, weighted by the time fraction *f* (see Eq. (2)). In addition, depletion times will in principle differ between resources. This could be captured by making the time fraction resource specific. In principle, we could have fitted resource-specific time fractions *f*_*i*_ directly from the community dynamics. However, this was not practical with the limited data we had access to without severe overfitting. Therefore, we simplified our model to use the same time fraction for all resources. This simplification can be partially justified because hCom2 was constructed by augmenting a simpler community (hCom1) with species from a large pool. This process filled all ecological niches that remained open in hCom1 and placed the most competitive strains in each niche. We have previously shown that in such mature communities, most of the time during each growth cycle is spent in the first temporal niche where all resources are present^16^. After this long first niche, all resources rapidly disappear one after another. Therefore, both the differences in the depletion times of the resources and the deviations of the approximate average abundances from the exact time averages are expected to be relatively small. This a posteriori justifies our simplifying assumptions.

Another approximation of our model was the use of coarse-grained resource clusters, which was necessary to avoid overfitting the data describing the growth of *∼* 60 species on nearly 300 resources. Our resource clustering strategy can be compared to previous work on consumer resource models (CRMs) for microbial communities of varying complexity. For low complexity communities, with up to 5 strains and resources, we and others have previously developed detailed CRMs^16,17,29,30^, where each resource has its own separate depletion time. These differences in resource depletion times could in principle be captured in our model with a resource-specific time fraction *f*_*i*_, but as explained above for the data we had for hCom2, this would lead to overfitting. At the intermediate complexity level, Ho et al. 2022^28^ developed a CRM for a simplified synthetic human gut microbiome with 15 strains. They used binary consumption fluxes of these strains to group resources into 2^15^ = 32, 768 binary groups based on the exact subset of species capable of consuming them. They then retained about 30 of the most abundant groups. This method is not scalable to a more complex community like the hCom2^11^ with 63 strains surviving in steady state. In fact, with binarized fluxes, there would be 2^63^ *∼* 10^19^ possible binary metabolite groups, and searching this prohibitively large space is computationally infeasible. Therefore, we resorted to a more traditional approach of clustering metabolites with similar consumption and production fluxes, but not necessarily identical binary consumption profiles. Our model can be easily adapted to work with individual metabolites if a sufficient number of experiments is available to estimate each *R*_*i*_ individually. One way to accomplish this is to run additional serial dilution experiments with inocula composed of subsets of strains of different diversity, in addition to the full diversity community where all strains are initially present.

Despite all the simplifications, our model performed reasonably well in predicting both the dynamics and the steady-state abundances of the strains in the serial dilution experiments of Ref.^31^. More than 50% of the residual error in the model predictions was due to biological replicate-to-replicate variability in the serial dilution abundance data (Fig. S3). Experimentally, this replicate-to-replicate variability is most likely a consequence of variation in the composition of the mega medium - a rich, undefined medium. Variability in the composition of the mega medium was also responsible for the day-to-day variation in consumption and production flux measurements observed in the experiments of Han et al. 2021^33^.

Using the model trained on the synthetic human gut microbiome hCom2, we performed three in-silico perturbation experiments to study the organizational properties of the community. From the leave-one-out experiment (Fig. 3b), we found that the intermediate abundance strains were the most sensitive to cooperative interactions (Fig. 3c). Strains of the genus *Bacteroides* were clearly overrepresented in this group (5 out of 7 labeled strains in Fig. 3c). This can be tentatively attributed to the fact that *Bacteroides* strains tend to be generalists^34,35^, making them likely recipients of cooperative cross-feeding interactions, which in turn places them in the intermediate abundance tier of a multi-level trophic community. It should be noted that the nutrient composition of the mega medium used in the in vitro experiments of Ref.^31^ is dramatically different from the polysaccharide-dominated environment of the human large intestine. Therefore, the trophic levels of *Bacteroides* strains in vivo^36,37^ are likely to be different from those observed in vitro.

The second in silico experiment we performed was to perturb metabolite clusters individually by increasing their concentration, *R*_*i*_, by a factor of 100 (Fig. 4). The effect of resources on strains can be classified as either direct or indirect. For direct effects, strain abundance increases in response to an increase in the concentration of a metabolite that it consumes. Strain abundances may also increase due to indirect interactions such as crossfeeding. Decreases in strain abundance in response to increases in the concentration of a single metabolite can only occur through indirect interactions such as resource competition. Our model captures both direct and indirect effects of perturbations in metabolite concentrations. This is reflected in increased (33%) or decreased (67%) strain abundances in response to 100-fold increases in concentrations of individual metabolites.

In the third in silico experiment, we designed and implemented a greedy algorithm to equalize steady-state strain abundances. This approach may be relevant for practical applications such as gut microbiome transplantation or supplementation therapy. Indeed, in a complex community with equalized abundances, all strains have a priori equal chances to survive and colonize the new environment, which in case of hCom2 is the gut of the microbiome transplant recipient^32^.

In conclusion, we have introduced and studied a simplified consumer resource model capable of predicting the dynamics of a complex community of strains in a serial dilution experiment. This model was tested on a defined synthetic human gut community consisting of a controlled collection of strains with known resource consumption and production fluxes. One of the potential future directions is to extend our model to other synthetic communities for which there is no metabolomics data to quantify consumption and production fluxes. One example of this with important practical applications is given by microbial strains isolated from the plant rhizosphere^38^. These strains were used to construct complex synthetic communities composed of 185 strains^14^ and 62 strains^39^ studied in *Arabidopsis thaliana* or 36 strains^13^ studied in Sorghum. The application of our model to these communities would require a reliable way to predict of consumption and production fluxes of individual strains directly from their genomes. While computational methods cannot fully replace dedicated metabolomics experiments, they can be used as a first-order approximation. Promising approaches include in silico reconstruction of mechanistic genome-scale metabolic models (see^40^ for a recent review) or “black box” machine learning algorithms to predict consumption and production of individual metabolites (see e.g.^41,42^).

## METHODS

### Data

We parameterized our model for a synthetic human gut microbiome hCom2^11^ using two published and publicly available datasets. The first is a metabolomics dataset comprising consumption and production fluxes of 178 strains, including all hCom2 strains, for 833 metabolites, generated using an integrated liquid chromatography-mass spectrometry (LC-MS) pipeline described in Han et al. 2021^33^. These strains were individually cultured in Mega Medium^43^-a rich, undefined medium known to support the growth of diverse bacteria. The culture supernatant was collected between mid-log and stationary phase for processing through the LC-MS pipeline. For each metabolite and strain, the consumption flux c _*α i*_ was calculated by subtracting from 1 the ratio of the concentrations of the metabolite *i* before and after batch growth of a single strain *γ* in the mega medium. Similarly, the total production flux 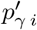 was calculated by subtracting 1 from the ratio of the metabolite *i* before and after batch growth of a single strain *γ* in the mega medium. The production flux *p*_*γ-i*_ used in Eq. 3b is calculated per unit biomass. It is given by the total production flux 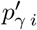 divided by the total biomass *N*_*γ*_(grown alone) of the strain *γ* at the end of the batch experiment. This biomass is computed as described below.

The second data set captures the dynamics of a synthetic human gut microbiome grown over 6 passages in a serial dilution experiment^31^. The set of 117 strains used in this experiment was modeled and extended from hCom2^11^. In one type of experiment, these strains were grown in a medium containing different types of beads that provided surfaces for bacterial attachment. This experiment was designed to mimic the spatial organization of the human gut microbiome. The second type of experiment was a control using only the liquid Mega Medium without beads. Our model assumes an equal dilution ratio of each strain and is only suitable for the control experiment without beads. In principle, it is possible to adapt our model to describe other experiments in Ref.^31^, where passage from one growth cycle to the next is achieved by transferring a single bead. However, it requires 63 new parameters that quantify the degree of adhesion of each of the strains to the beads. It is not computationally feasible to fit these parameters without additional experiments, so we limited our study to no-bead control experiments. Cultures were grown in Mega Medium^43^ for three days and then passaged with the dilution factor *D* = 15, 000. Already after three serial passages the community was observed to reach the steady state^31^. Therefore, we limited our model to describe the community dynamics during the first three passages. Each serial passaging experiment was run in three biological replicates, with each biological replicate after each passage sequenced in three technical replicates. For our analysis, we averaged the abundances from the technical replicates. Our model was necessarily limited to include only the 63 strains from Ref.^31^ for which we had consumption and production fluxes from Ref.^33^. However, these strains accounted for *∼* 90% of the total abundance of all strains surviving in the steady state and thus provided a reasonably good approximation to the full community studied in Ref.^31^. The names of these strains, along with the abbreviations we assigned to them, are listed in Table S1.

### Clustering metabolites

The number of strains (63) used in our model is much smaller than the initial number of individual metabolites (292) in the Mega Medium. To avoid severe overfitting, we clustered the metabolites using the consumption fluxes. The clustering procedure gave us 10 non-singleton and 88 singleton metabolite clusters. These 98 clusters were used for all analyses in the paper. The names of the metabolites in each metabolite cluster are given in Table. S2. Some metabolite names are present more than once in the metabolomics data from Han et al. 2021^33^. For these, we preserved these repeats by adding numerical suffixes.

For clustering we first binarized the consumption fluxes *c*_*α-i*_ with a threshold of 0.3, and used only those metabolites consumed by more than 5 strains. These metabolites were grouped into 10 non-singleton clusters using hierarchical agglomerative clustering with Euclidean distance as the metric and Ward’s linkage method. The remaining metabolites consumed by less than 5 strains were used as singleton clusters. In total, we obtained 98 metabolite clusters.

To get *c*_*α-i*_ for a non-singleton cluster *i*, we first took the average over all metabolites *j* in this cluster: 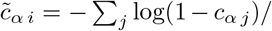 (# of metabolites in the cluster) Then, *c*_*α-i*_ for the cluster was obtained by the inverse transformation 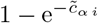.

Similarly, for the total metabolite 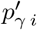 for the non-singleton cluster *i*, we took the average 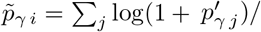 (# of metabolites in the cluster). Then, the production flux for the cluster was obtained by the inverse transformation 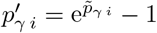.

### Estimation of *R*_*i*_ for metabolite clusters

*R*_*i*_ were estimated from Eq. (3) using *c*_*α-i*_, *p*_*γ-i*_, *N*_*α*_(end of cycle) and *f* as inputs. The strain abundances were used only for the passages 1 and 2. These two passages were the most dynamic for hCom2^31^. Passages 1 and 2 were used as *N*_*α*_(end of cycle 1) and *N*_*α*_(end of cycle 2), respectively. Eq. (3) was applied to each biological replicate. This resulted in a system of linear equations in *R*_*i*_, which was solved using non-negative least squares (NNLS)^44^ to obtain an estimate of non-negative *R*_*i*_. This NNLS problem was solved using the nnls function from the optimize subpackage of the SciPy Python package^45^.

The production flux *p*_*γ-i*_ used in Eq. (3b) is given by the total metabolite production flux 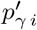 derived from the metabolomics experiment^33^ normalized by the biomass of this strain *N*_*γ*_(grown alone) in the end of the batch experiment: 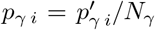 (grown alone). In practice, we do not know these abundances but can estimate them using the mass conservation: *N*_*γ*_(grown alone) = *i R*_*i*_ *c*_*γ-i*_, which in turn depends on *R*_*i*_. We used an iterative approach to jointly solve Eq. (3) for *R*_*i*_ and *N*_*γ*_(grown alone). First, we assigned *N*_*γ*_(grown alone) = 1*/n*_*s*_. Next, we inferred *R*_*i*_ and used this value to update our estimate for *N*_*γ*_(grown alone) using the aforementioned approximation. The updated *N*_*γ*_(grown alone) was used to obtain the next estimate for *R*_*i*_. This iterative scheme was repeated for 100 iterations. The values of *N*_*γ*_(grown alone) and *R*_*i*_ obtained at the end of the procedure were used for downstream analyses.

### Predicting strain abundances from the model

With *R*_*i*_ obtained from the NNLS fit as described above, we used Eq. (3) to predict the strain abundances forward in time. This was done starting with the experimentally measured strain abundances in the inoculum (averaged over two biological replicates) as *N*_*α*_(end of cycle 0). Given *R*_*i*_, *c*_*α-i*_, *p*_*γ-i*_, *N*_*α*_(end of cycle 0) and *f* (see the next section on how we fitted *f*), Eqs. (3) for each *α* fully define *N*_*α*_(end of cycle 1) (the number of equations is equal to the number of unknowns). We solved this system of nonlinear equations iteratively. First, *N*_*α*_(end of cycle 1) were set equal to *N*_*α*_(end of cycle 0) on the right-hand side (RHS) of Eq. (3). The RHS was then used to calculate the new estimate for *N*_*α*_(end of cycle 1) on the left-hand side (LHS). This new estimate was again substituted into the RHS to obtain an updated estimate for *N*_*α*_(end of cycle 1). This was continued for 100 iterations to obtain the final estimate for *N*_*α*_(end of cycle 1).

There were two biological replicates for the inoculum, which were not matched to the three biological replicates of serial passage experiments. Therefore, to predict strain abundances, we averaged the two inoculum abundances and used that as the starting point. As a result the model generates only one prediction for all passages, whereas the observed data had three different biological replicates. We took the geometric average (on log_10_ scale) of the three biological replicates to compare against the model prediction. The variation in the three biological replicates is shown as horizontal error bars in Fig. 2.

### Estimation of the time fraction *f*

To fit the time fraction parameter *f* for hCom2, we estimated *R*_*i*_ for linearly spaced values of *f* in the interval [0.3, 1] as shown in Fig. S1. For each estimated *R*_*i*_, we made a prediction for the steady-state strain abundances reached at the third passage starting from the inoculum. The predicted abundances were compared to the observed abundances. As mentioned before, we took the geometric average (on log_10_ scale) of the three biological replicates to compare against the model prediction. The value of *f* that gave the best agreement with the observed data was selected. The quality of the agreement was quantified by the Pearson’s correlation coefficient between the logarithms of the predicted and observed abundances averaged over three biological replicates. Using this approach we found a sharp peak around *f* = 0.9 for hCom2 (Fig. S1). Therefore, *f* = 0.9 was used in our model throughout this study.

### Estimate of replicate-to-replicate variability in experimentally observed abundances

For each passage, except the inoculum, three biological replicates were provided for the abundance of each strain^31^. To estimate the replicate-to-replicate variability at each passage, for a given pair of replicates, Root Mean Square Error (RMSE) was calculated using log_10_ of strain abundances. This was repeated for the 3 possible replicate pairs. The average RMSE for the 3 pairs of biological replicates was used as an estimate of the replicate-to-replicate variability.

### Estimation of the cumulative RMSE over steady-state abundances

To calculate the cumulative RMSE for either the predictive performance of the model or the replicate-to-replicate variability in Fig. S3a-c, the strains were sorted in an increasing order of observed steady-state abundances (averaged over biological replicates). We then considered a set of different abundance thresholds (evenly spaced on the log_10_ scale). For each threshold, all strains above that threshold were retained and the others were dropped. The RMSE of log_10_ of the strain abundances was calculated for the subset of strains exceeding a given threshold.

### Average competition and the cross-feeding scores between strains

For each pair of bacterial strains, we calculate a ‘competition score’. This score shows how much they compete for the same food resources, which in this case are metabolites or metabolite clusters. We only look at metabolites that are present in the system (those with a non-zero concentration *R*_*i*_). To make this score easy to understand, we rescale it to a scale from 0 to 1. This is done by dividing the number of shared metabolites by the total number of metabolites consumed by the perturbed strain. We average this score across all strain pairs to get an overall competition score.

Similarly, we calculate a ‘cross-feeding score’. This score measures how many metabolites produced by one strain are consumed by another strain. This score is also normalized to be between 0 and 1 by dividing it by the number of metabolites consumed by the perturbed strain.

### Greedy algorithm to equalize strain abundances

At each iteration of our algorithm, we consider the effect of increasing or decreasing the concentration of a single metabolite *R*_*i*_ on the strain abundances and choose a perturbation that brings them closest to uniformity. When increasing the concentration, we simply multiply *R*_*i*_ by 10. Metabolites initially absent in the medium are assigned a nominal very low concentration *R*_*i*_ = 10^*−*8^. Since we do not allow the concentration of any metabolite to fall below its unperturbed value in the Mega Medium, we have implemented a special rule for decreasing the concentration, summarized in the following equation:

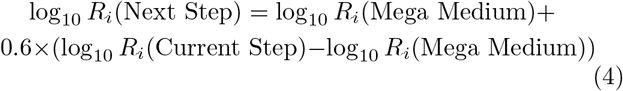

This rule ensures that the recipe discovered by our algorithm can be implemented experimentally by supplementing the Mega Medium with prescribed concentrations of selected metabolites. Indeed, there is no practical way to remove individual metabolites from a complex undefined medium such as the Mega Medium. At each iteration, for each *R*_*i*_, a fair coin toss decides whether the concentration is increased or decreased, which adds stochasticity to the algorithm. The greedy step of our algorithm is repeated up to 500 times or until no further improvements can be made (local uniformity optimum).

The algorithm was repeated 10 times to obtain 10 possible perturbed *R*_*i*_’s, which were then processed as described in the Results section.

## Acknowledgements

We thank Veronika Dubinkina and Akshit Goyal for useful discussions.

## Supplementary Figures and Tables

**FIG. S1.**
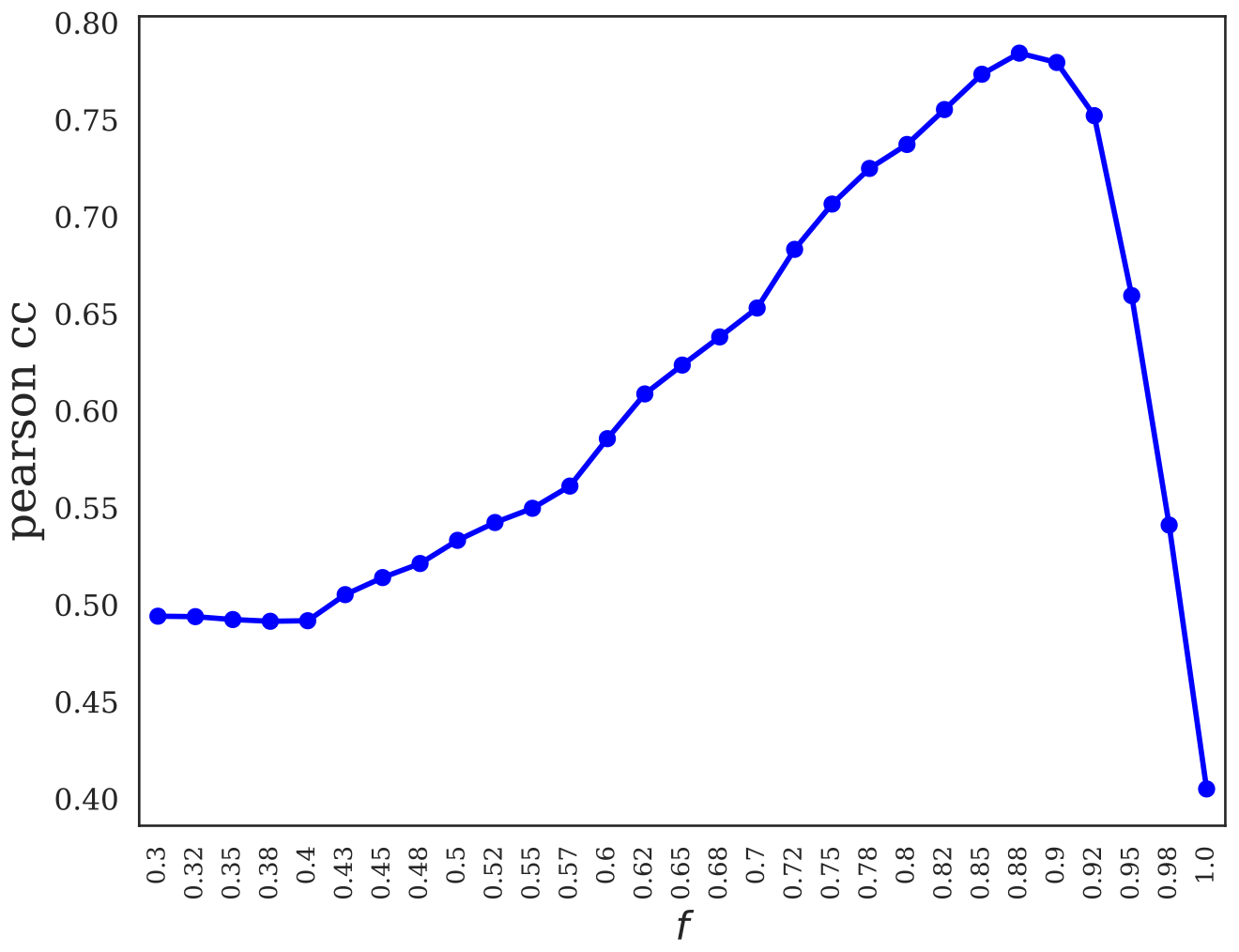
Estimating the time fraction *f* for hCom2. The time fraction *f* was estimated by inferring *R*_*i*_ from our model for linearly spaced values of *f* in the interval [0.3, 1]. For each estimated *R*_*i*_, we made a prediction of the steady-state strain abundances starting from the inoculum using our model. Pearson’s correlation coefficients between predicted and observed abundances for different values of *f* are plotted.

**TABLE S1.**
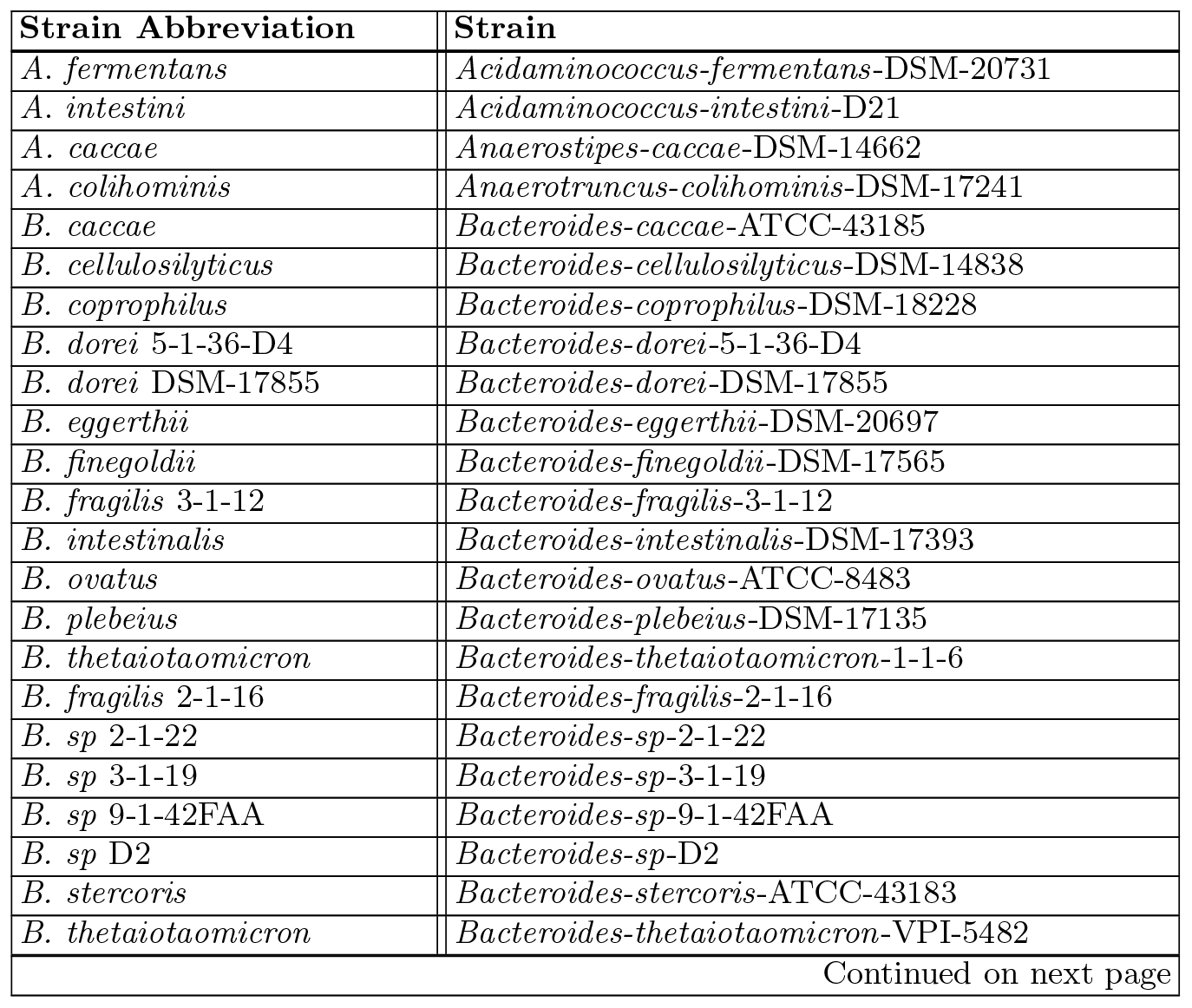
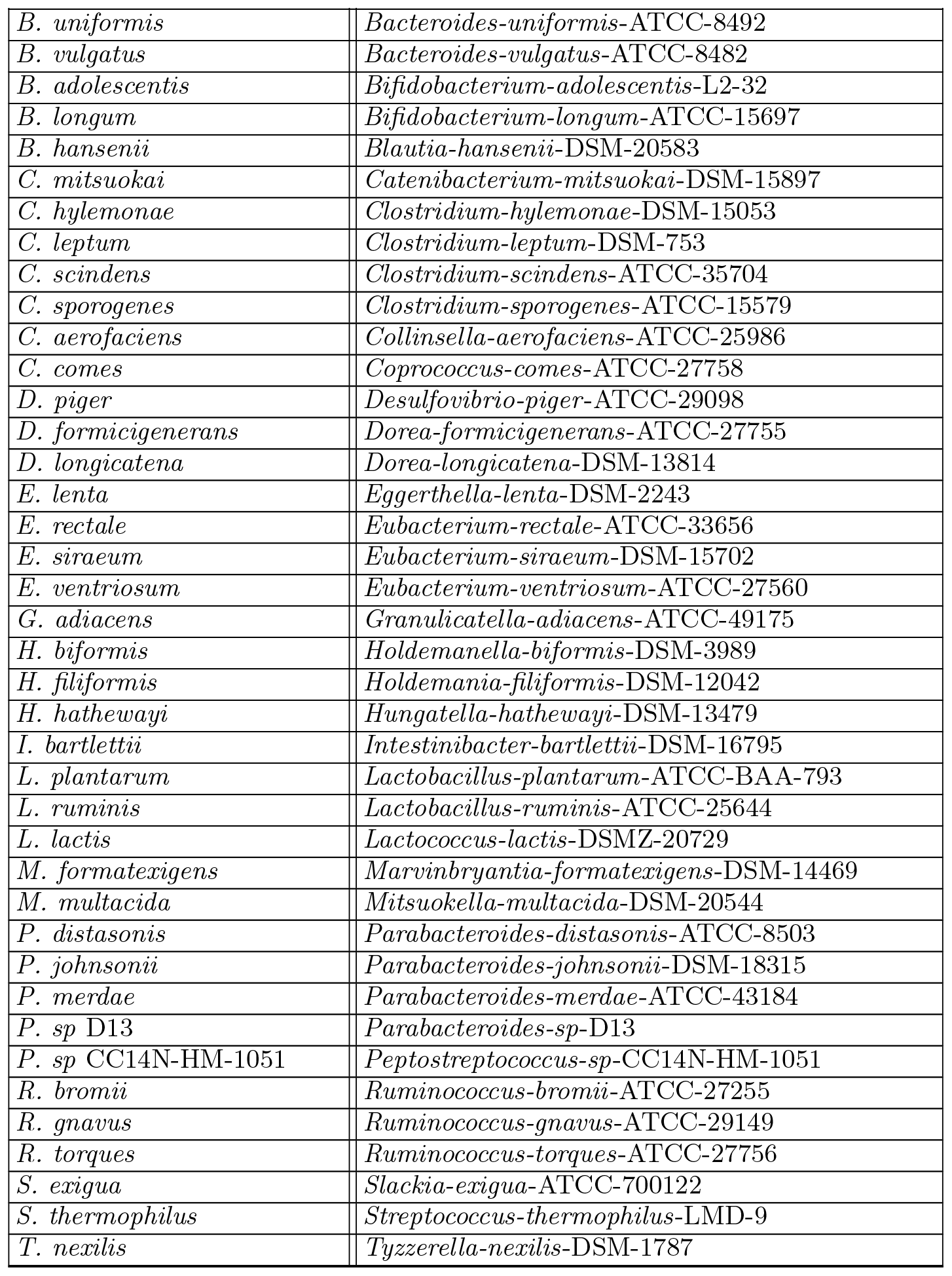
Abbreviations for strain names.

**TABLE S2.**
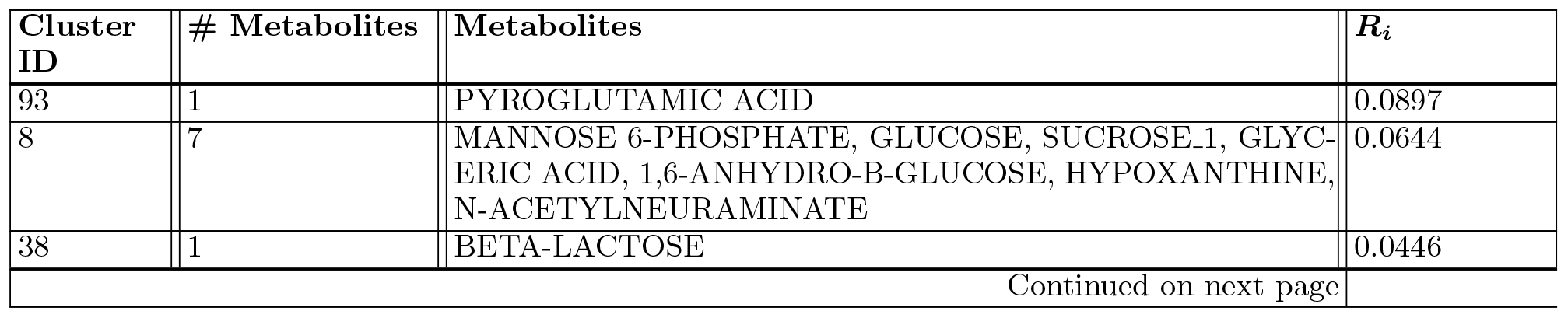
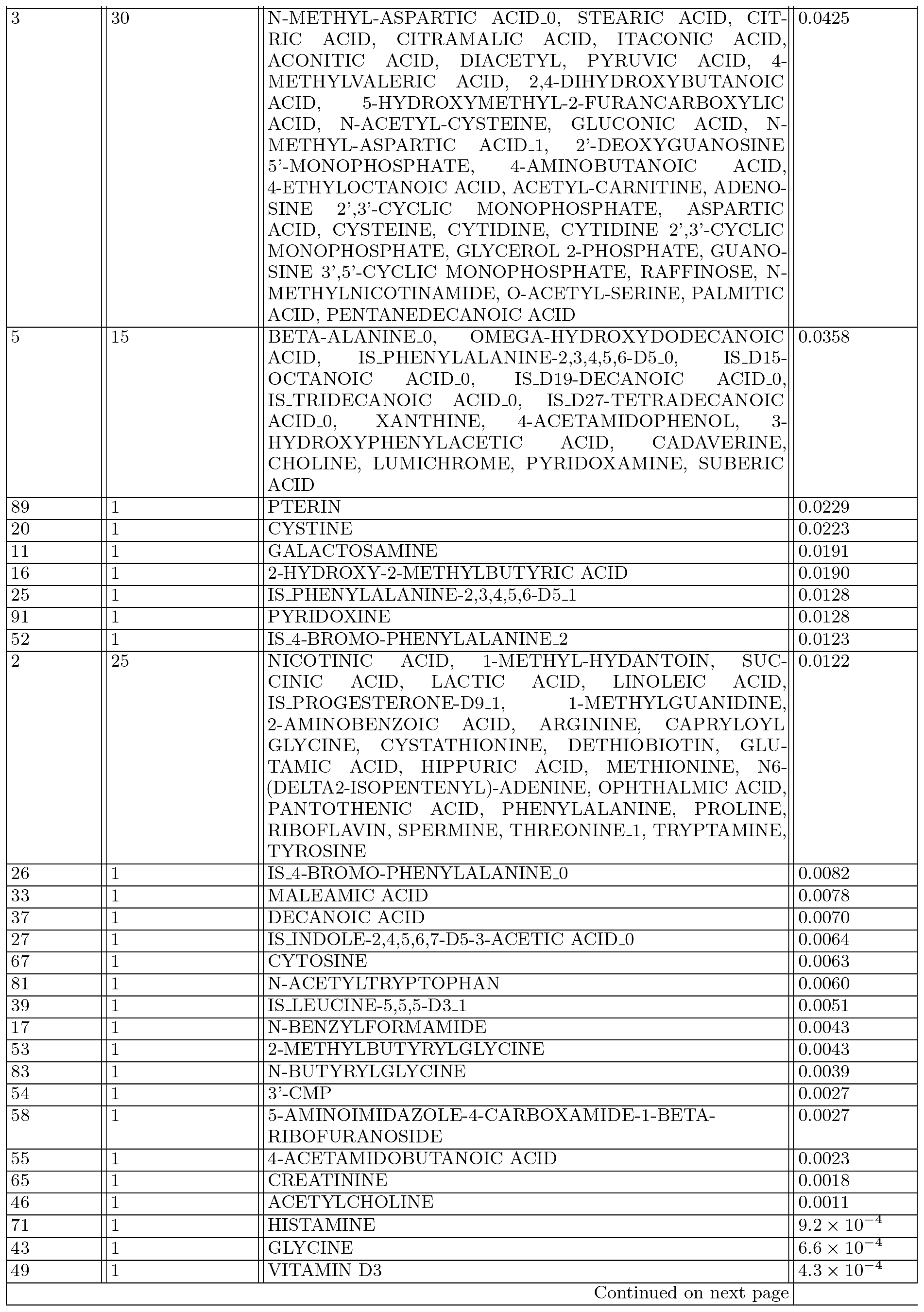
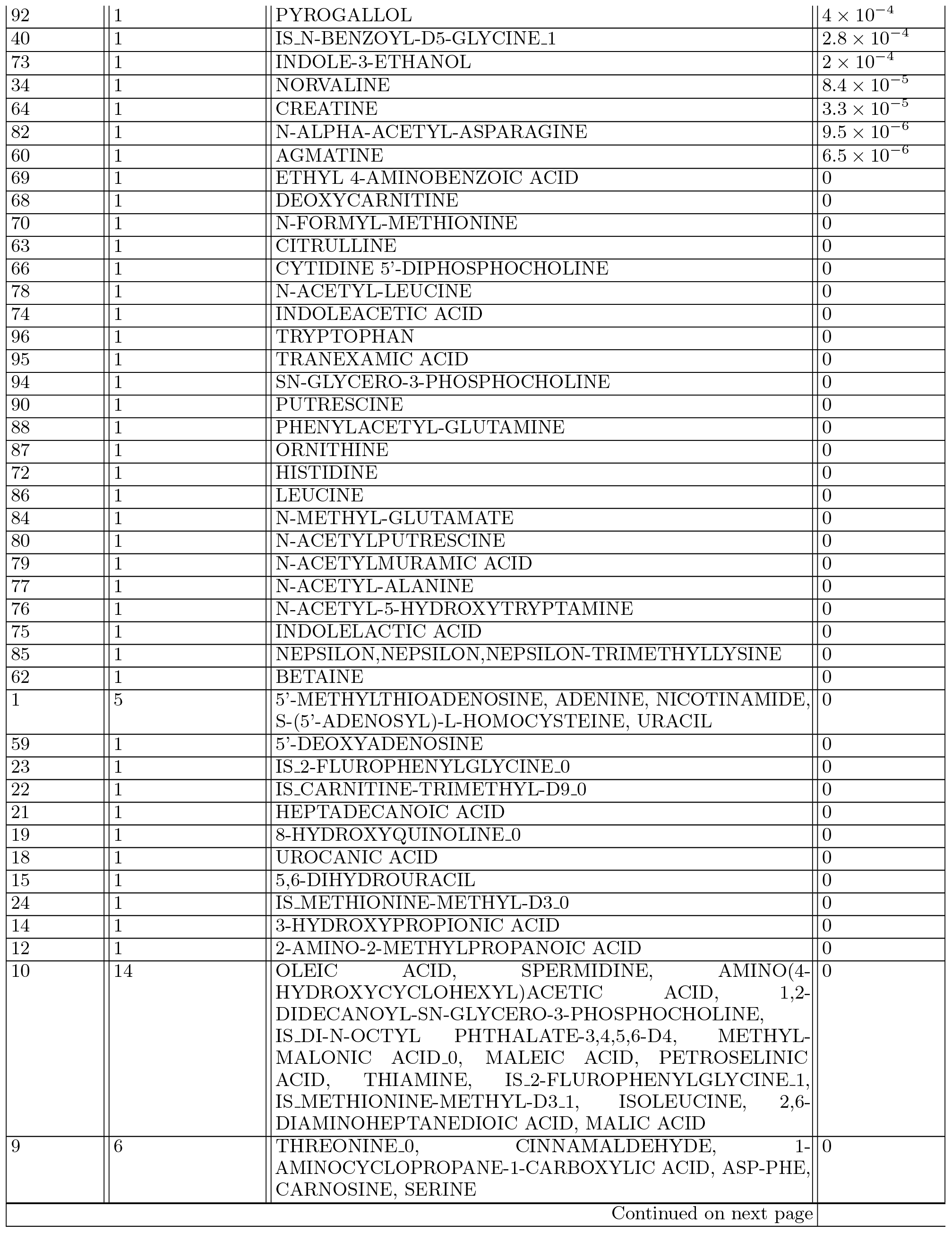
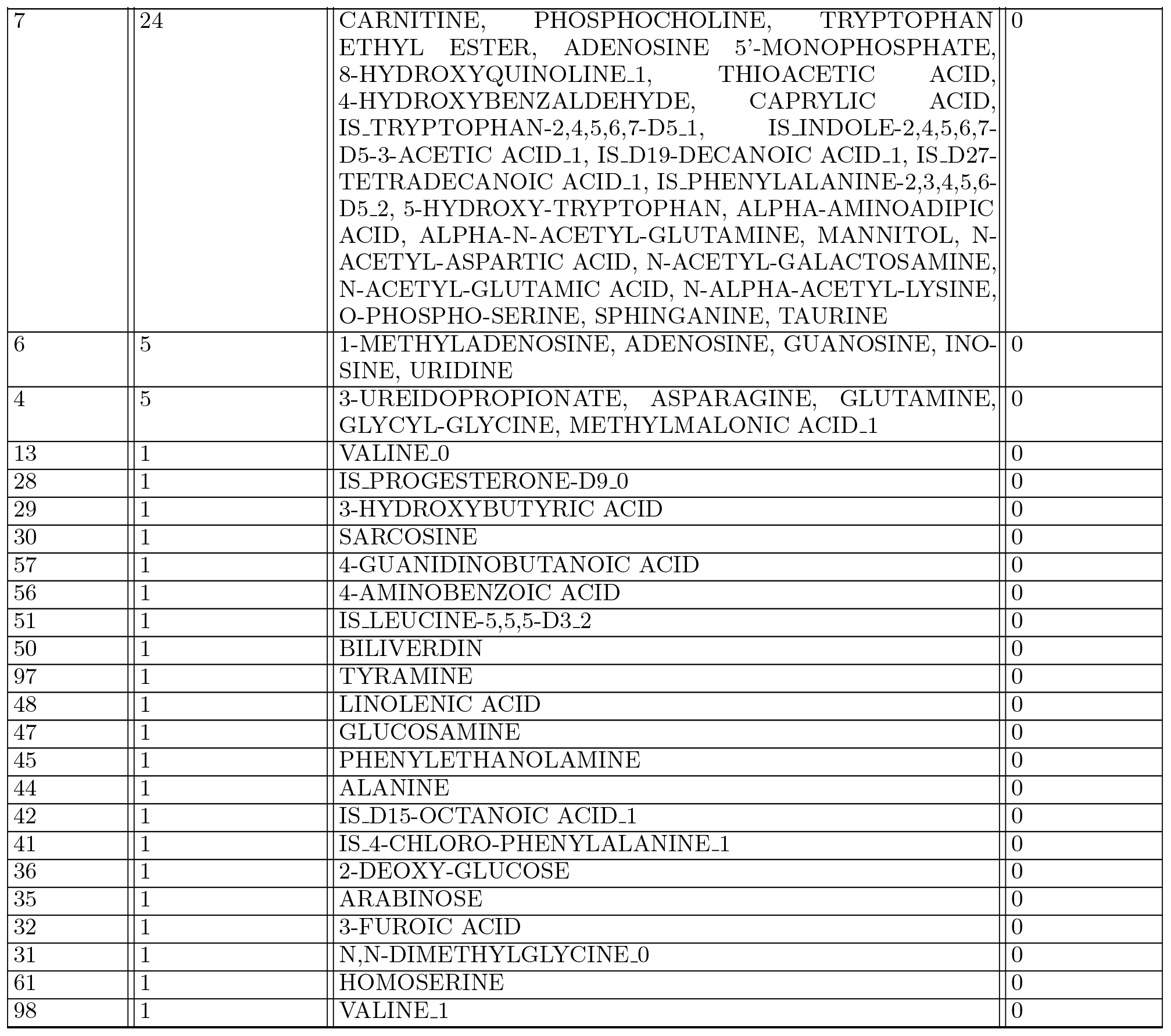
Metabolite clusters sorted in decreasing order of *R*_*i*_ for hCom2.

| Cluster ID | # Metabolites | Metabolites | $R_i$ |
| --- | --- | --- | --- |
| 93 | 1 | PYROGLUTAMIC ACID | 0.0897 |
| 8 | 7 | MANNOSE 6-PHOSPHATE, GLUCOSE, SUCROSE_1, GLYC-ERIC ACID, 1,6-ANHYDRO-B-GLUCOSE, HYPOXANTHINE, N-ACETYLNEURAMINATE | 0.0644 |
| 38 | 1 | BETA-LACTOSE | 0.0446 |

TABLE S2 – continued from previous page
| Cluster ID | # Metabolites | Metabolites | $R_i$ |
| --- | --- | --- | --- |
| 3 | 30 | N-METHYL-ASPARTIC ACID_0, STEARIC ACID, CITRIC ACID, CITRAMALIC ACID, ITACONIC ACID, ACONITIC ACID, DIACETYL, PYRUVIC ACID, 4-METHYLVALERIC ACID, 2,4-DIHYDROXYBUTANOIC ACID, 5-HYDROXYMETHYL-2-FURANCARBOXYLIC ACID, N-ACETYL-CYSTEINE, GLUCONIC ACID, N-METHYL-ASPARTIC ACID_1, 2'-DEOXYGUANOSINE 5'-MONOPHOSPHATE, 4-AMINOBUTANOIC ACID, 4-ETHYLOCTANOIC ACID, ACETYL-CARNITINE, ADENOSINE 2',3'-CYCLIC MONOPHOSPHATE, ASPARTIC ACID, CYSTEINE, CYTIDINE, CYTIDINE 2',3'-CYCLIC MONOPHOSPHATE, GLYCEROL 2-PHOSPHATE, GUANOSINE 3',5'-CYCLIC MONOPHOSPHATE, RAFFINOSE, N-METHYLNICOTINAMIDE, O-ACETYL-SERINE, PALMITIC ACID, PENTANEDECANOIC ACID | 0.0425 |
| 5 | 15 | BETA-ALANINE_0, OMEGA-HYDROXYDODECANOIC ACID, IS-PHENYLALANINE-2,3,4,5,6-D5_0, IS_D15-OCTANOIC ACID_0, IS_D19-DECANOIC ACID_0, IS_TRIDECANOIC ACID_0, IS_D27-TETRADECANOIC ACID_0, XANTHINE, 4-ACETAMIDOPHENOL, 3-HYDROXYPHENYLACETIC ACID, CADAVERINE, CHOLINE, LUMICHRONE, PYRIDOXAMINE, SUBERIC ACID | 0.0358 |
| 89 | 1 | PTERIN | 0.0229 |
| 20 | 1 | CYSTINE | 0.0223 |
| 11 | 1 | GALACTOSAMINE | 0.0191 |
| 16 | 1 | 2-HYDROXY-2-METHYLBUTYRIC ACID | 0.0190 |
| 25 | 1 | IS-PHENYLALANINE-2,3,4,5,6-D5_1 | 0.0128 |
| 91 | 1 | PYRIDOXINE | 0.0128 |
| 52 | 1 | IS.4-BROMO-PHENYLALANINE_2 | 0.0123 |
| 2 | 25 | NICOTINIC ACID, 1-METHYL-HYDANTOIN, SUCINIC ACID, LACTIC ACID, LINOLEIC ACID, IS-PROGESTERONE-D9_1, 1-METHYLGUANIDINE, 2-AMINOBENZOIC ACID, ARGININE, CAPRYLOYL GLYCINE, CYSTATHIONINE, DETHIOBIOTIN, GLUTAMIC ACID, HIPPURIC ACID, METHIONINE, N6-(DELTA2-ISOPENTENYL)-ADENINE, OPTHALMIC ACID, PANTOTHENIC ACID, PHENYLALANINE, PROLINE, RIBOFLAVIN, SPERMINE, THREONINE_1, TRYPTAMINE, TYROSINE | 0.0122 |
| 26 | 1 | IS.4-BROMO-PHENYLALANINE_0 | 0.0082 |
| 33 | 1 | MALEAMIC ACID | 0.0078 |
| 37 | 1 | DECANOIC ACID | 0.0070 |
| 27 | 1 | IS.INDOLE-2,4,5,6,7-D5-3-ACETIC ACID_0 | 0.0064 |
| 67 | 1 | CYTOSINE | 0.0063 |
| 81 | 1 | N-ACETYLTRYPTOPHAN | 0.0060 |
| 39 | 1 | IS.LEUCINE-5,5,5-D3_1 | 0.0051 |
| 17 | 1 | N-BENZYLFORMAMIDE | 0.0043 |
| 53 | 1 | 2-METHYLBUTYRYLGLYCINE | 0.0043 |
| 83 | 1 | N-BUTYRYLGLYCINE | 0.0039 |
| 54 | 1 | 3'-CMP | 0.0027 |
| 58 | 1 | 5-AMINOIMIDAZOLE-4-CARBOXAMIDE-1-BETA-RIBOFURANOSIDE | 0.0027 |
| 55 | 1 | 4-ACETAMIDOBUTANOIC ACID | 0.0023 |
| 65 | 1 | CREATININE | 0.0018 |
| 46 | 1 | ACETYLCHOLINE | 0.0011 |
| 71 | 1 | HISTAMINE | $9.2 \times 10^{-4}$ |
| 43 | 1 | GLYCINE | $6.6 \times 10^{-4}$ |
| 49 | 1 | VITAMIN D3 | $4.3 \times 10^{-4}$ |

TABLE S2 – continued from previous page
| Cluster ID | # Metabolites | Metabolites | $R_i$ |
| --- | --- | --- | --- |
| 92 | 1 | PYROGALLOL | $4 \times 10^{-4}$ |
| 40 | 1 | IS_N-BENZOYL-D5-GLYCINE_1 | $2.8 \times 10^{-4}$ |
| 73 | 1 | INDOLE-3-ETHANOL | $2 \times 10^{-4}$ |
| 34 | 1 | NORVALINE | $8.4 \times 10^{-5}$ |
| 64 | 1 | CREATINE | $3.3 \times 10^{-5}$ |
| 82 | 1 | N-ALPHA-ACETYL-ASPARAGINE | $9.5 \times 10^{-6}$ |
| 60 | 1 | AGMATINE | $6.5 \times 10^{-6}$ |
| 69 | 1 | ETHYL 4-AMINOBENZOIC ACID | 0 |
| 68 | 1 | DEOXYCARNITINE | 0 |
| 70 | 1 | N-FORMYL-METHIONINE | 0 |
| 63 | 1 | CITRULLINE | 0 |
| 66 | 1 | CYTIDINE 5'-DIPHOSPHOCHOLINE | 0 |
| 78 | 1 | N-ACETYL-LEUCINE | 0 |
| 74 | 1 | INDOLEACETIC ACID | 0 |
| 96 | 1 | TRYPTOPHAN | 0 |
| 95 | 1 | TRANEXAMIC ACID | 0 |
| 94 | 1 | SN-GLYCERO-3-PHOSPHOCHOLINE | 0 |
| 90 | 1 | PUTRESCINE | 0 |
| 88 | 1 | PHENYLACETYL-GLUTAMINE | 0 |
| 87 | 1 | ORNITHINE | 0 |
| 72 | 1 | HISTIDINE | 0 |
| 86 | 1 | LEUCINE | 0 |
| 84 | 1 | N-METHYL-GLUTAMATE | 0 |
| 80 | 1 | N-ACETYLPUTRESCINE | 0 |
| 79 | 1 | N-ACETYLMURAMIC ACID | 0 |
| 77 | 1 | N-ACETYL-ALANINE | 0 |
| 76 | 1 | N-ACETYL-5-HYDROXYTRYPTAMINE | 0 |
| 75 | 1 | INDOLELACTIC ACID | 0 |
| 85 | 1 | NEPSILON,NEPSILON,NEPSILON-TRIMETHYLLYSINE | 0 |
| 62 | 1 | BETAINE | 0 |
| 1 | 5 | 5'-METHYLTHIOADENOSINE, ADENINE, NICOTINAMIDE, S-(5'-ADENOSYL)-L-HOMOCYSTEINE, URACIL | 0 |
| 59 | 1 | 5'-DEOXYADENOSINE | 0 |
| 23 | 1 | IS_2-FLUOROPHENYLGLYCINE_0 | 0 |
| 22 | 1 | IS_CARNITINE-TRIMETHYL-D9_0 | 0 |
| 21 | 1 | HEPTADECANOIC ACID | 0 |
| 19 | 1 | 8-HYDROXYQUINOLINE_0 | 0 |
| 18 | 1 | UROCANIC ACID | 0 |
| 15 | 1 | 5,6-DIHYDROURACIL | 0 |
| 24 | 1 | IS_METHIONINE-METHYL-D3_0 | 0 |
| 14 | 1 | 3-HYDROXYPROPIONIC ACID | 0 |
| 12 | 1 | 2-AMINO-2-METHYLPROPANOIC ACID | 0 |
| 10 | 14 | OLEIC ACID, SPERMIDINE, AMINO(4-HYDROXYCYCLOHEXYL)ACETIC ACID, 1,2-DIDECANOYL-SN-GLYCERO-3-PHOSPHOCHOLINE, IS_DI-N-OCTYL PHTHALATE-3,4,5,6-D4, METHYL-MALONIC ACID_0, MALEIC ACID, PETROSELINIC ACID, THIAMINE, IS_2-FLUOROPHENYLGLYCINE_1, IS_METHIONINE-METHYL-D3_1, ISOLEUCINE, 2,6-DIAMINOHEPTANEDIOIC ACID, MALIC ACID | 0 |
| 9 | 6 | THREONINE_0, CINNAMALDEHYDE, 1-AMINOCYCLOPROPANE-1-CARBOXYLIC ACID, ASP-PHE, CARNOSINE, SERINE | 0 |

TABLE S2 – continued from previous page
| Cluster ID | # Metabolites | Metabolites | $R_i$ |
| --- | --- | --- | --- |
| 7 | 24 | CARNITINE, PHOSPHOCHOLINE, TRYPTOPHAN ETHYL ESTER, ADENOSINE 5'-MONOPHOSPHATE, 8-HYDROXYQUINOLINE_1, THIOACETIC ACID, 4-HYDROXYBENZALDEHYDE, CAPRYLIC ACID, IS.TRYPTOPHAN-2,4,5,6,7-D5_1, IS.INDOLE-2,4,5,6,7-D5-3-ACETIC ACID_1, IS.D19-DECANOIC ACID_1, IS.D27-TETRADECANOIC ACID_1, IS.PHENYLALANINE-2,3,4,5,6-D5_2, 5-HYDROXY-TRYPTOPHAN, ALPHA-AMINOADIPIC ACID, ALPHA-N-ACETYL-GLUTAMINE, MANNITOL, N-ACETYL-ASPARTIC ACID, N-ACETYL-GALACTOSAMINE, N-ACETYL-GLUTAMIC ACID, N-ALPHA-ACETYL-LYSINE, O-PHOSPHO-SERINE, SPHINGANINE, TAURINE | 0 |
| 6 | 5 | 1-METHYLADENOSINE, ADENOSINE, GUANOSINE, INOSINE, URIDINE | 0 |
| 4 | 5 | 3-UREIDOPROPIONATE, ASPARAGINE, GLUTAMINE, GLYCYL-GLYCINE, METHYLMALONIC ACID_1 | 0 |
| 13 | 1 | VALINE_0 | 0 |
| 28 | 1 | IS.PROGESTERONE-D9_0 | 0 |
| 29 | 1 | 3-HYDROXYBUTYRIC ACID | 0 |
| 30 | 1 | SARCOSINE | 0 |
| 57 | 1 | 4-GUANIDINOBUTANOIC ACID | 0 |
| 56 | 1 | 4-AMINOBENZOIC ACID | 0 |
| 51 | 1 | IS.LEUCINE-5,5,5-D3_2 | 0 |
| 50 | 1 | BILIVERDIN | 0 |
| 97 | 1 | TYRAMINE | 0 |
| 48 | 1 | LINOLENIC ACID | 0 |
| 47 | 1 | GLUCOSAMINE | 0 |
| 45 | 1 | PHENYLETHANOLAMINE | 0 |
| 44 | 1 | ALANINE | 0 |
| 42 | 1 | IS.D15-OCTANOIC ACID_1 | 0 |
| 41 | 1 | IS.4-CHLORO-PHENYLALANINE_1 | 0 |
| 36 | 1 | 2-DEOXY-GLUCOSE | 0 |
| 35 | 1 | ARABINOSE | 0 |
| 32 | 1 | 3-FUROIC ACID | 0 |
| 31 | 1 | N,N-DIMETHYLGLYCINE_0 | 0 |
| 61 | 1 | HOMOSERINE | 0 |
| 98 | 1 | VALINE_1 | 0 |

**FIG. S2.**
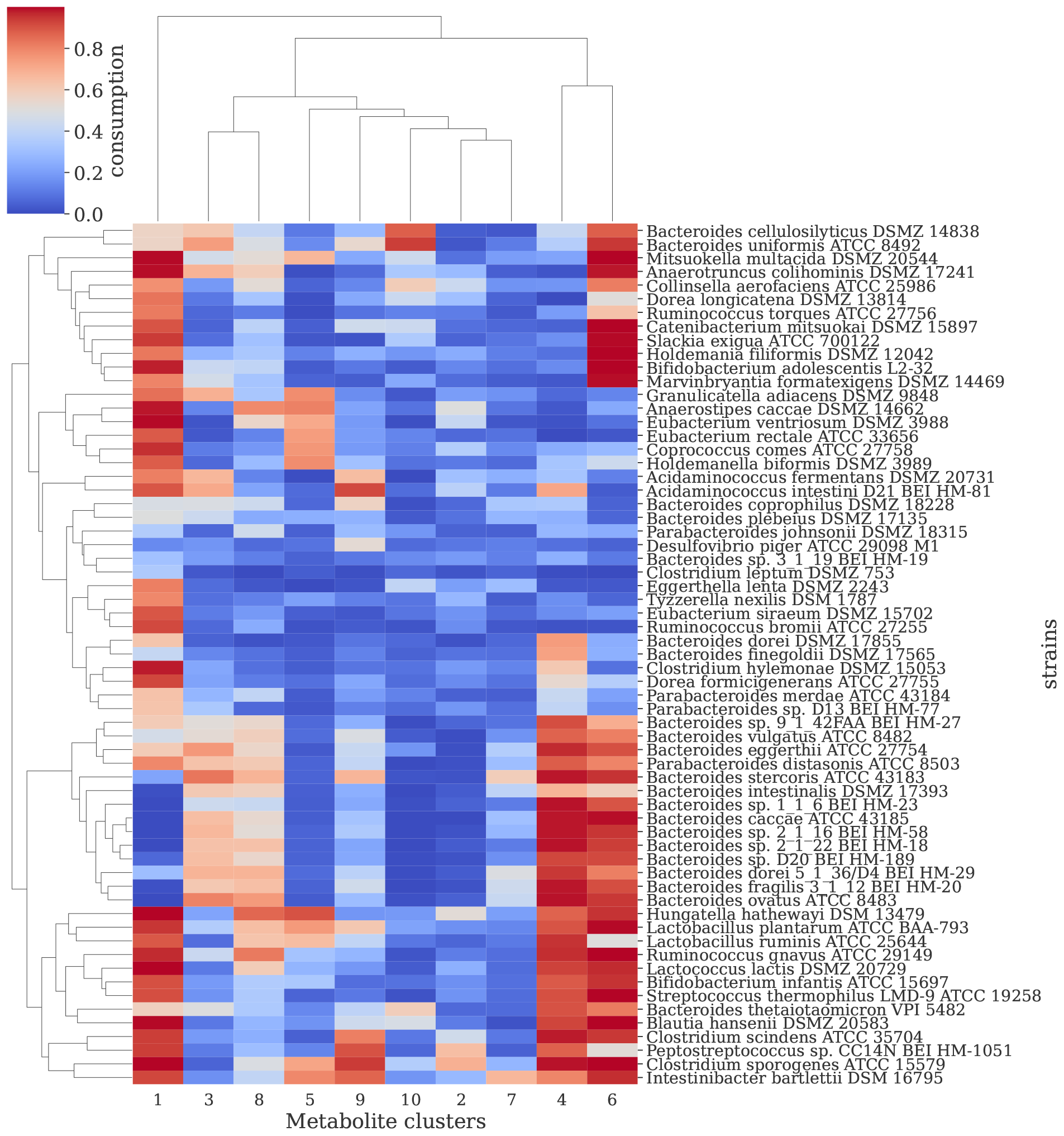
Hierarchically clustered heatmap of the consumption fluxes for the 10 non-singleton metabolite clusters for hCom2. Columns and rows represent the metabolite clusters and the strains, respectively.

**FIG. S3.**
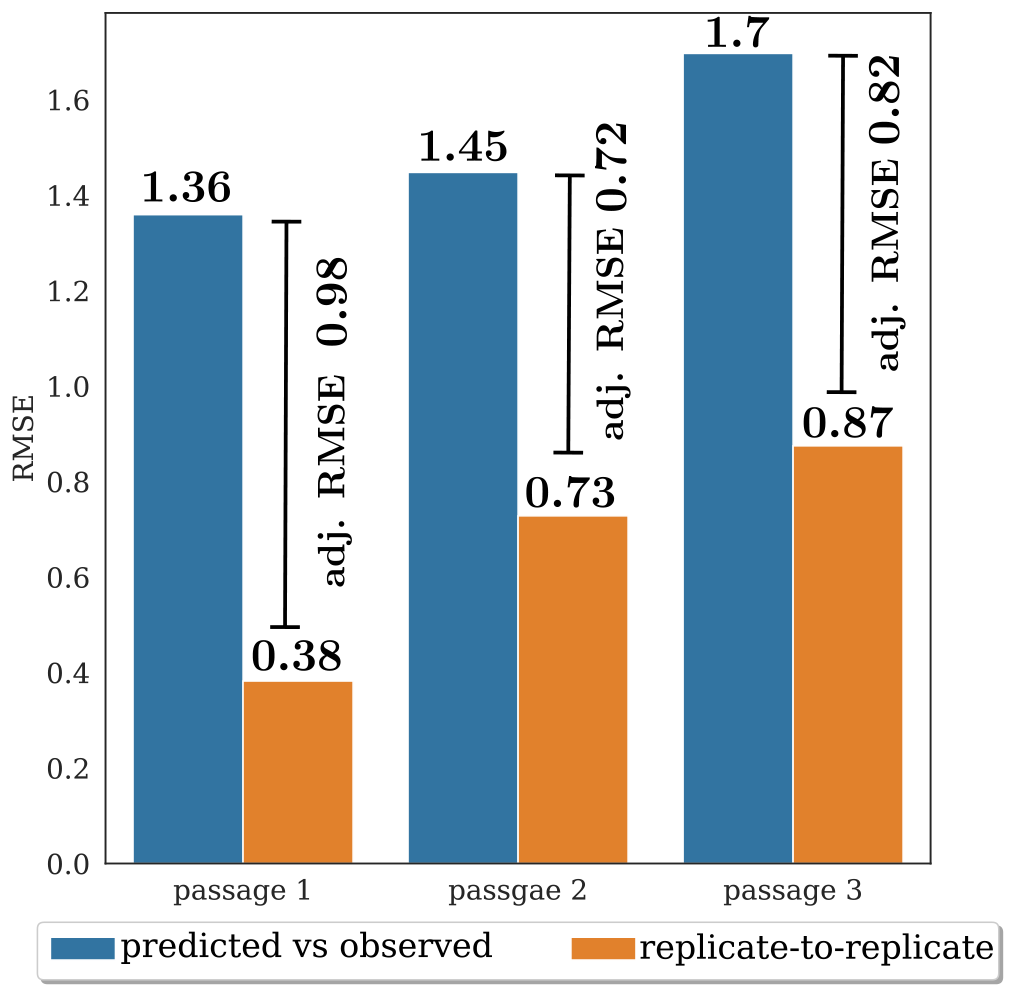
Biological replicate-to-replicate variability explains more than half of the residual error in our model predictions for hCom2. Biological replicate-to-replicate variability at each passage was estimated as the root mean squared error (RMSE) between the strain abundances for the biological replicates. Model performance at each passage was estimated as the RMSE between predicted and observed strain abundances. The adjusted RMSE was then calculated as the difference between the RMSE for model performance and the RMSE for biological replicate-to-replicate variability.

**FIG. S4.**
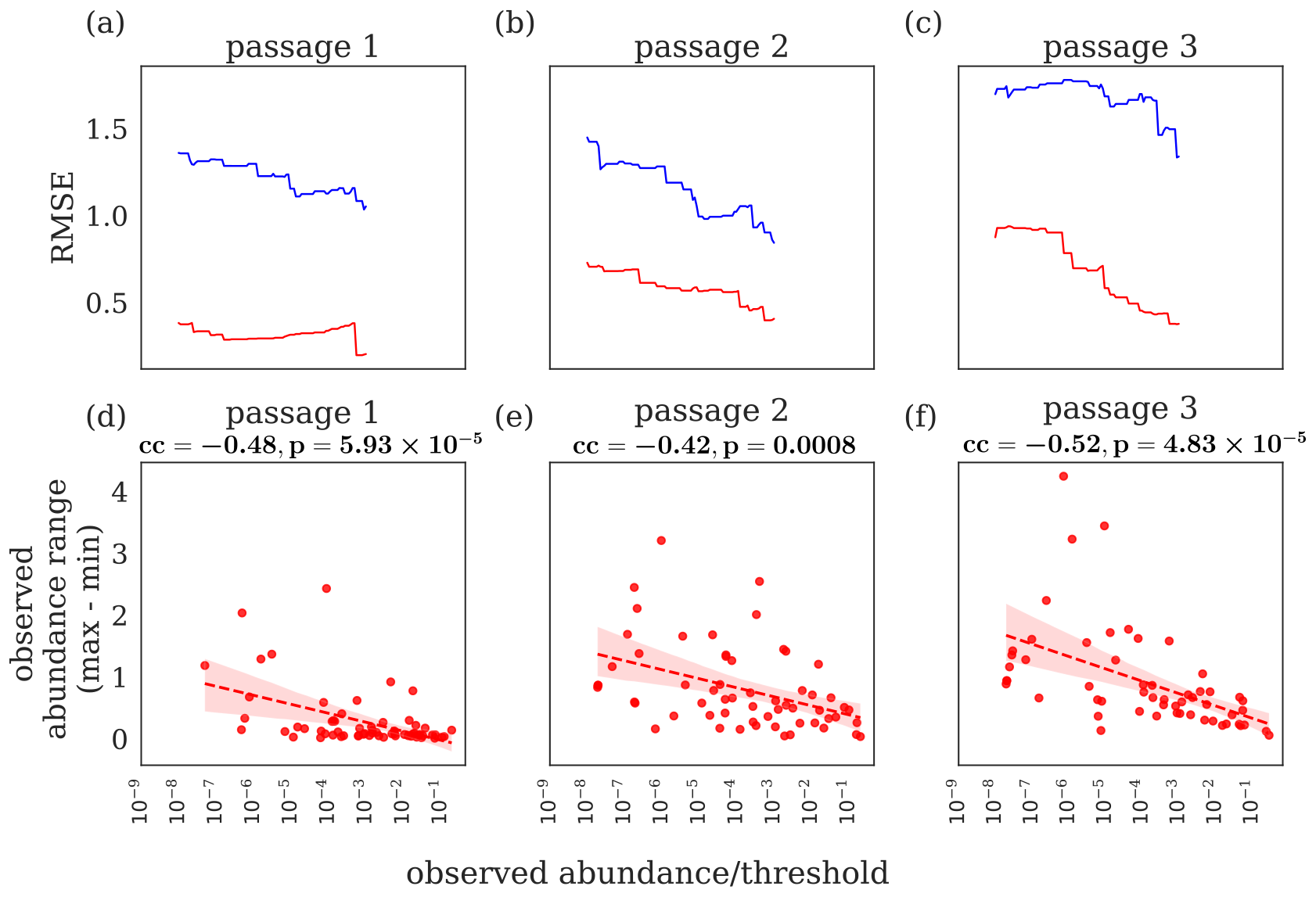
Biological replicate-to-replicate variability and model performance are negatively correlated with strain abundances in hCom2. **a)-c)** Cumulative RMSE is plotted as a function of average observed strain abundances at steady state for model performance (blue curve) and biological replicate-to-replicate variability (red curve). First, the strains were sorted in increasing order of average observed steady-state abundances. For each value of observed steady-state abundance used as a threshold, the cumulative RMSE at each passage was calculated for a subset of strains with strain abundances at that passage greater than the threshold (details are given in the main Methods section). **d)-f)** The width of the error bars (maximum minus minimum) from Fig. 2 are plotted as a function of observed strain abundance for different passages. Each point on the scatterplot represents a single strain. Pearon’s correlation coefficient (cc) (along with the p-value) is given for the different passages. The linear regression fit to the scatterplot is also shown as a dashed line, with the 95% confidence interval shown as a shaded area around the linear fit.

**FIG. S5.**
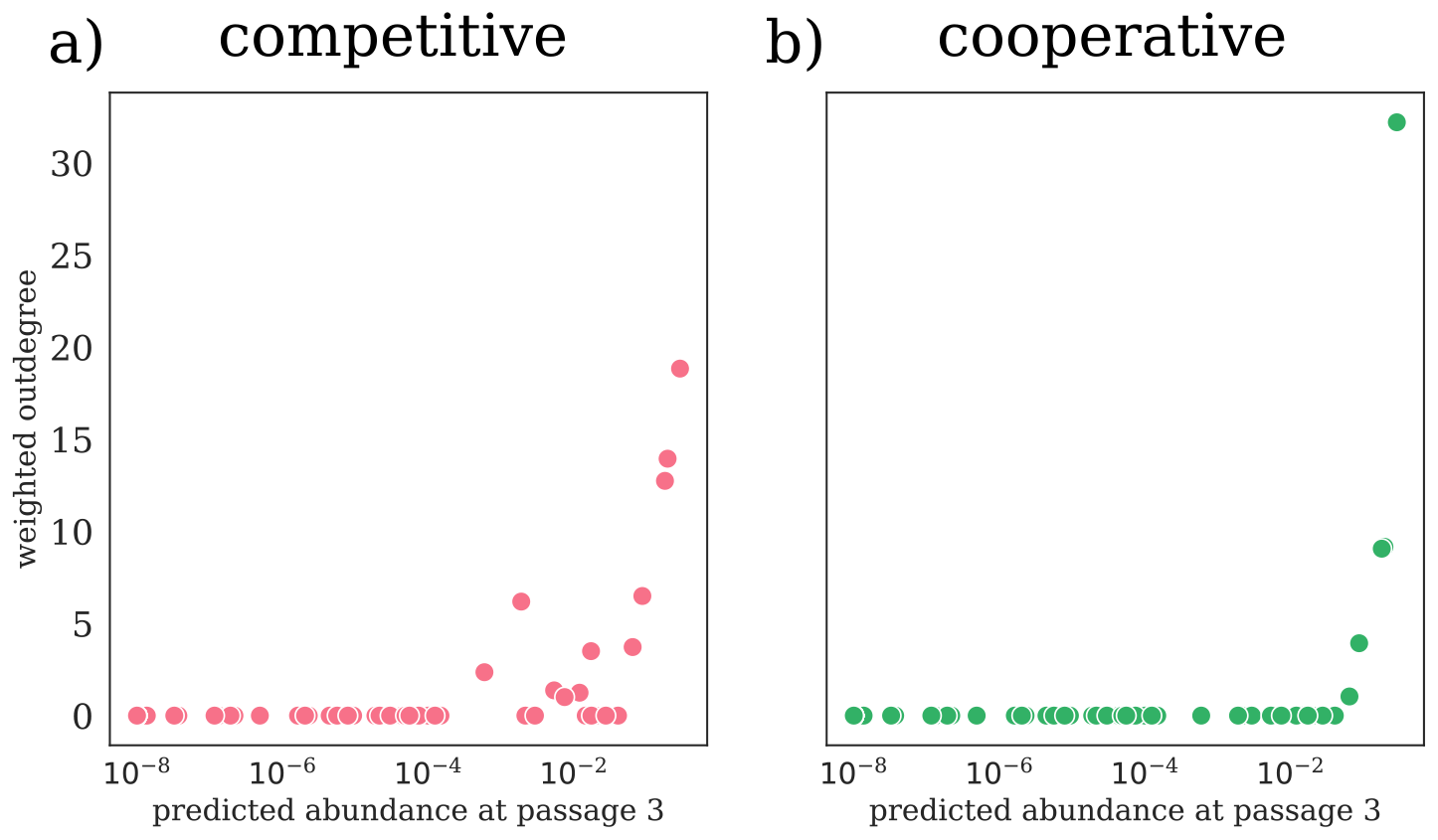
Weighted strain out-degree versus steady-state abundance for hCom2. Total weighted out-degree restricted to competitive (red, **a)**) and cooperative (red, **b)**) edges in the strain-strain interaction network for all the strains plotted against predicted strain abundances at steady state (passage 3) for the unperturbed hCom2 community.

**FIG. S6.**
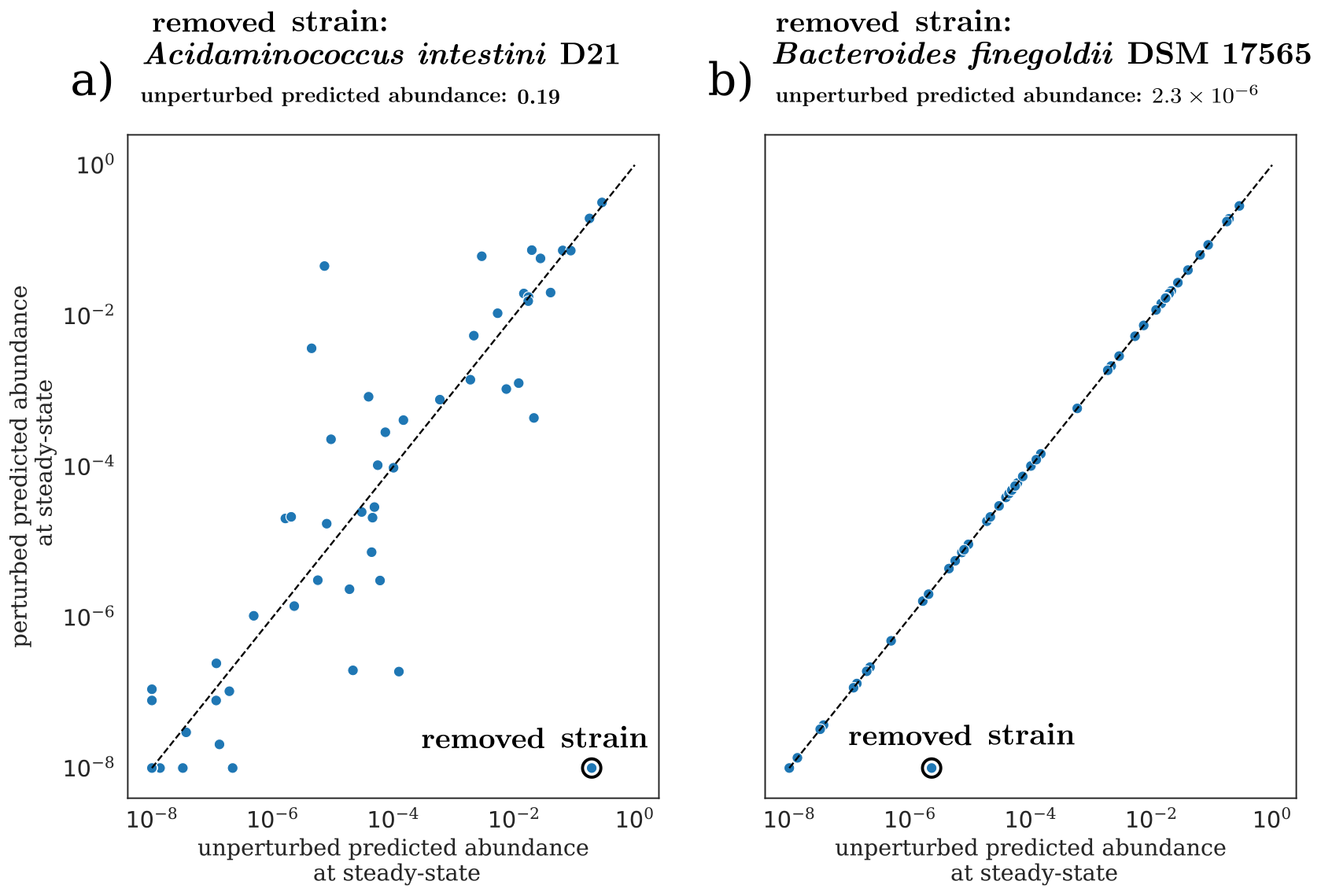
Perturbations from leave-one-experiment are non-trivial and cannot be explained by re-normalization for relative abundances for hCom2. **a)** Removal of the highest abundance strain from hCom2 causes steady-state abundances for multiple strains to change by different factors, which cannot be explained by re-normalization of perturbed abundances. **b)** Removal of a low abundance strain does not change steady-state abundances for the hCom2 community.

**FIG. S7.**
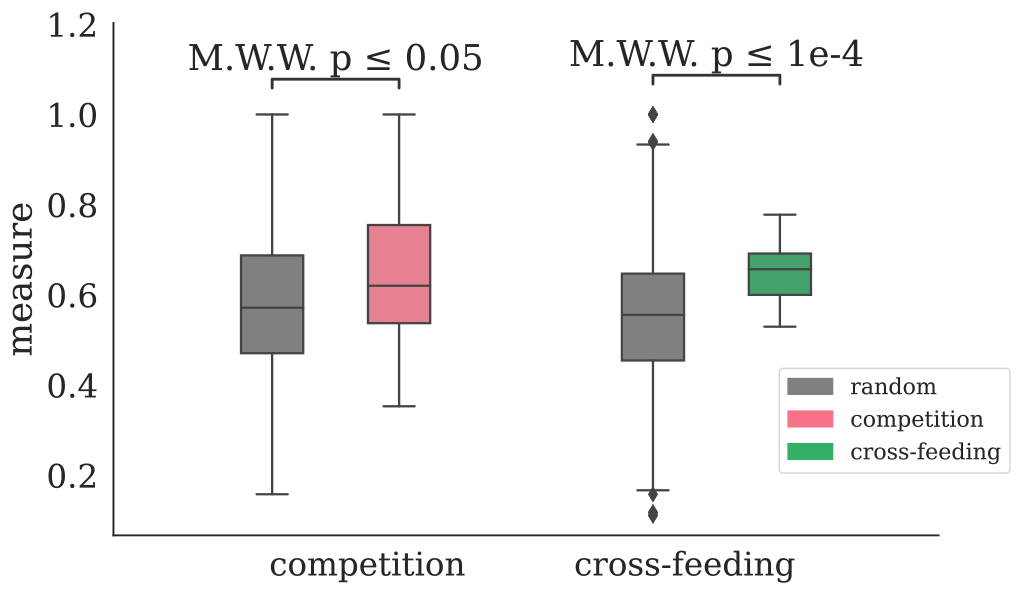
Competition and cross-feeding underlie competitive and cooperative edges for hCom2. (Left) Comparison of competition scores for competitive edges (red) against random edges. Non-interacting and cooperative edges (green) are defined as random in this context. We obtained a mean competition score of 0.65 for competitive edges vs. 0.58 for non-interacting or cooperative interaction edges (*p* = 0.02, two-tailed Mann-Whitney Wilcoxon test). (Right)Comparison of cross-feeding scores for cooperative edges (green) against random edges. Non-interacting and competitive edges (red) are defined as random in this context. We obtained a mean cross-feeding score of 0.64 for cooperative edges vs. 0.55 for non-interacting or competitive interaction edges (*p* = 9.7 *×* 10^*−*5^, two-tailed Mann-Whitney Wilcoxon test).

**FIG. S8.**
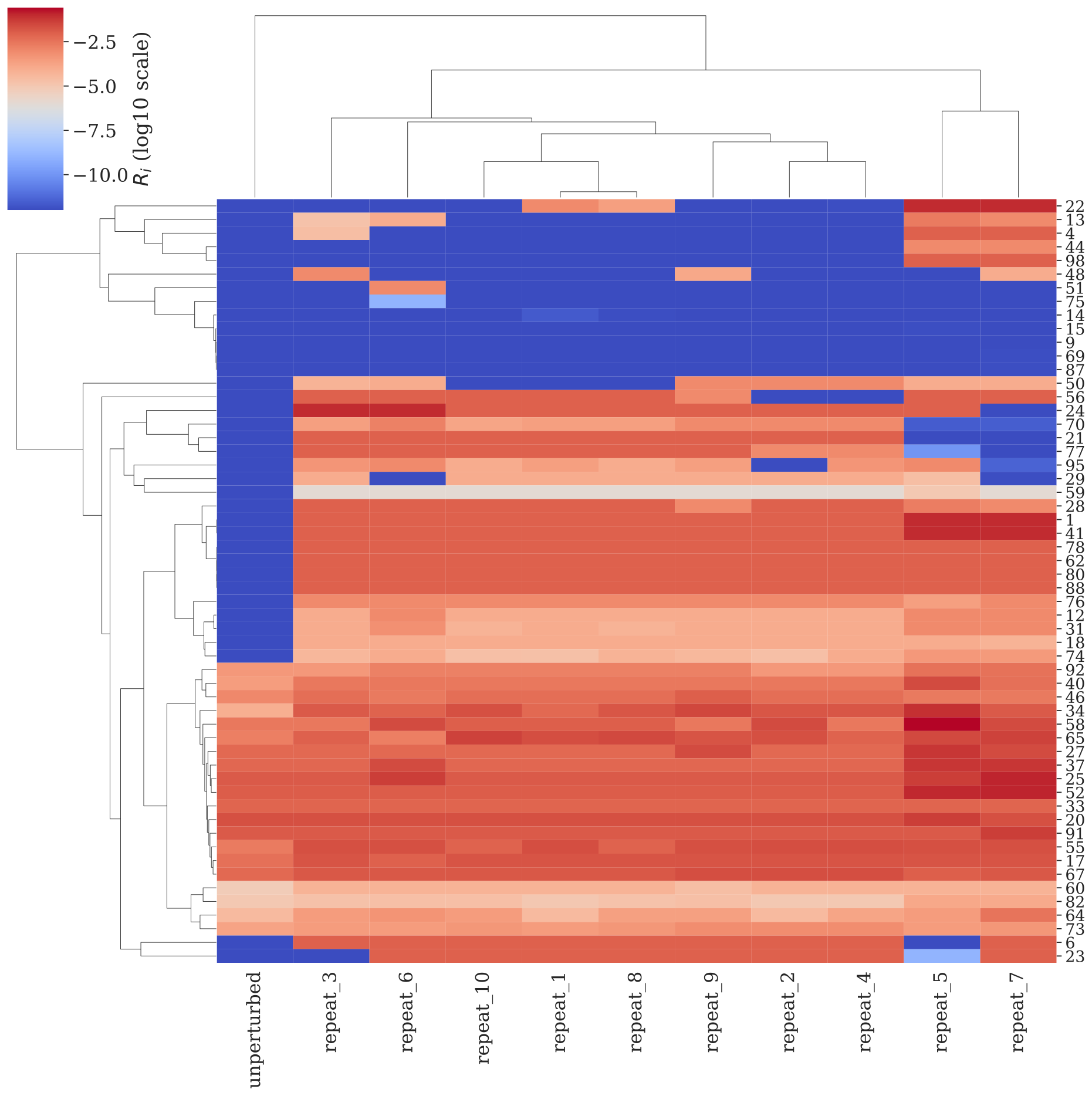
Hierarchically clustered heatmap of the 10 perturbed *R*_*i*_ solutions generated by our greedy algorithm for equalising strain abundances at steady-state for hCom2. The columns and rows represent the different solutions and metabolite clusters, respectively.

